# Gut Microbiota-derived Adenosine Determines the Efficacy of Electroconvulsive Therapy for Depression

**DOI:** 10.1101/2025.11.13.688184

**Authors:** Yinglei Wang, Mengyao Dai, Dongdong Yang, Siyu Fan, Yanzhang Li, Hanqing Zhao, Zhiling Ye, Qian Xu, Yiping Dou, Zili Han, Yang Ji, Yue Wu, Kai Wang, Shu Zhu, Chuan Huang, Yanghua Tian

**Affiliations:** State Key Laboratory of Immune Response and Immunotherapy, School of Basic Medical Sciences, Division of Life Sciences and Medicine, University of Science and Technology of China, Hefei, China; Institute of Health and Medicine, Hefei Comprehensive National Science Center, Hefei, China; Department of Neurology, The First Affiliated Hospital of Anhui Medical University, Hefei, China; Department of Neurology, The Third Affiliated Hospital of Anhui Medical University, Hefei, China; Department of Neurology, The Second Affiliated Hospital of Anhui Medical University, Hefei, China; Department of Psychology and Sleep Medicine, The Second Affiliated Hospital of Anhui Medical University, Hefei, China; The School of Mental Health and Psychological Sciences, Anhui Medical University, Hefei, China

## Abstract

Electroconvulsive therapy (ECT) is a final procedure for major depressive disorders (MDD). However, the effectiveness of ECT in treating MDD patients varies. Here we identified gut microbiota as a potential restraining factor of ECT efficacy. By using a 5-day electroconvulsive shock (ECS) in depression mouse model of chronic restraint stress (CRS), it was observed that the antidepressant effect of ECS was suppressed after antibiotics (ABX)-induced microbiota dysbiosis, which also resulted in elevated c-Fos expression of CRH neurons in paraventricular nucleus of hypothalamus (PVN) and increased serum corticosterone levels after ECS. Metabolomic analysis revealed the critical involvement of microbiota associated purine metabolism, and the supplement of gut microbiota-derived adenosine was able to recover the antidepressant effect of ECS in ABX-treated CRS mouse model. Based on the neural tracing from colon to brain and the immunofluorescence staining, it was surmised a gut-brain axis originated from Adora1-expressing cells in colon mediate this biological process. Furthermore, adenosine supplement alleviated the further deterioration of post-ECS memory loss and impairment of synaptic plasticity induced by ABX-treatment as well. Our findings suggest a potential role of microbial adenosine metabolism in ECT efficacy determination and side effect protection, highlights potential application of adenosine supplement in antidepressant therapies of those depression patients with microbiota dysbiosis.

## Introduction

Since first introduced as a treatment for psychiatric disorders in the 1930s, electroconvulsive therapy (ECT) has been proven to have strong positive effects on severe, treatment-resistant mental health conditions^[1, 2]^. Studies have indicated that the use of ECT results in a decreased risk of suicide^[3]^, improved functional outcomes and quality of life^[4]^, and decreased rates of rehospitalization in patients with depression^[5]^. Yet, the efficacy of ECT is still limited. A meta-analysis of studies of ECT for MDD showed a significantly lowered overall remission rate in patients with treatment resistant depression^[6]^, and a clinical trial also reported that only 25.32% (39/154) of MDD patients showed fast improvement after ECT^[7]^. In addition, ECT still causes several adverse effects, including amnesia, headache, muscle aches, nausea and fatigue^[8]^. Most patients have mild or moderate cognitive side effects, which usually resolve within days to weeks after the completion of the ECT course^[9–11]^. Retrograde amnesia is the most common persistent adverse effect of ECT^[1]^. Although retrograde amnesia often retrieves during the first few months after ECT, for many patients, recovery is incomplete, with prolonged amnesia regarding events that occurred close to the time of treatment^[12]^.

Different ECT procedures and different conditions of MDD patients can affect ECT efficacy and adverse effects^[13–18]^. Right unilateral and bifrontal placement may be selected to reduce the burden of side effects, whereas bilateral placement may be selected if the right unilateral or bifrontal positions are unlikely to be effective^[14, 15, 18]^. Age influences ECT efficacy as well: old patients response ECT faster than younger ones and require less sessions to achieve response and remission^[16, 17]^. Some case reports have indicated connections between microbiota and ECT^[19, 20]^. A recent clinical study on patients with severe or treatment-resistant depression has reported a significant difference of alpha diversity of oral microbiota between ECT responders and non-responders^[21]^, indicating an association between microbiota and ECT efficacy.

ECS mimics ECT in rodents to study mechanisms underlying the antidepressant effects and adverse effects of ECT^[22]^. ECS treatment has been reported to involve cortical network remodeling, in which controlled electrical stimulation induces cortical spreading depolarization^[23]^, a transient depression of cortical activity that is thought to reset dysfunctional neural circuits, and in models with impaired synaptic architecture, ECS also restores synaptic density and improves behavioral outcomes^[24]^, suggesting that the remodeling of brain network activity plays an important role in the effects of ECS. In addition, a study found that ECS significantly upregulated the expression of immediate early gene c-Fos and synaptic protein Narp in hippocampus, and promoted neurogenesis and dendritic shaping, suggesting that synaptic plasticity is crucial for the therapeutic effect^[25]^. Only a few articles have explored the antidepressant mechanism of ECS from a metabolic perspective. It has been reported that ECS can gradually restore mitochondrial bioenergetics after repeated treatment in a chronic stress model, it can improve oxygen consumption rates and neuronal ATP levels, reduce lactate accumulation, and increase pyruvate levels. These changes indicate that mitochondrial oxidative phosphorylation is enhanced, and brain energy metabolism is improved after ECS treatment^[26]^.

Adenosine, a neuromodulator derived from ATP metabolism, plays a critical role in regulating synaptic transmission, neuronal excitability, and neuroinflammation. Its role in the central nervous system (CNS) has gained increasing attention for its involvement in mood regulation and as a potential target in the treatment of major depressive disorder (MDD). Adenosine exerts its neuromodulatory effects through four G-protein-coupled receptors: A1, A2A, A2B, and A3. Among these, A1 and A2A receptors are most prominently expressed in tissues and are directly implicated in depressive behavior modulation. Activation of A1 receptors generally leads to inhibitory effects on neurotransmission, reducing neuronal excitability and promoting sleep-like states^[27]^. Astrocyte-derived adenosine activates A1 receptors to induce an antidepressant response in animal models. Mice lacking A1 receptors do not exhibit this antidepressant-like behavior, highlighting the interaction between adenosine and A1 receptors in antidepressant effect^[28]^. Emerging research also indicates that adenosine’s antidepressant properties may be partially mediated by modulation of the gut microbiota. A study demonstrated that oral administration of adenosine ameliorated depressive-like behaviors and corrected gut dysbiosis in a mouse model of social defeat stress. Parallel findings of reduced levels of short-chain fatty acids (SCFAs) in adolescents with depression further supported correlations between serum SCFA, adenosine levels, and microbial diversity^[29]^.

In this study, we identified gut microbiota as a potential restraining factor of ECT efficacy. We performed a 5-day ECS after ABX-induced dysbiosis in CRS mouse model and assessed behavior profiles and neuronal activities after ECS treatment. A fecal metabolomic analysis based on LC-MS figured out how gut microbiome interact with ECS efficacy and further validated the function of specific metabolite based on the metabolomic analysis. Our findings may provide a new vision in improving ECT efficacy and other antidepressant therapies.

## Results

### Gut microbiota influences ECS antidepressant effect in CRS mice

According to the analysis of clinical data from MDD patients, the decrease in Hamilton depression (HAMD)^[30]^ scores after ECT showed a significant positive correlation with the Shannon entropy of alpha-diversity of their fecal samples collected before ECT, which indicate the diversity of the gut microbiota. (Fig. 1A, Fig. S1A-C)^[31, 32]^. Gut microbiota is a key factor in maintaining intestinal homeostasis and affects multiple physiological systems of the host, including the nervous system^[33]^, this result suggests an association of ECT efficacy and gut homeostasis. To investigate whether gut microbiota influences ECT efficacy, an electroconvulsive shock (ECS) model in mouse was generated to mimic ECT, and different periods of ECS were tested to determine the duration of treatment. After 3-weeks chronic restraint stress (CRS) modeling, each group was treated with a 1/3/5/8-day ECS once a day (Fig. S1D-E). After ECS treatment, the depressive-like behaviors were assessed among all the groups, and decreased depressive-like behaviors were observed, presenting as increased preference in the sucrose preference test (SPT) without altered total consumption, and decreased immobility in the tail suspension test (TST) and the forced swimming test (FST) (Fig. S1F-J). The antidepressive-like behaviors reached the peak when treated with a 5-day ECS. Based on that, we used 5-day ECS model for following experiments. Furthermore, an antibiotic mix (ABX)-induced gut microbiota dysbiosis model was generated in mouse, combined with CRS and ECS to assess their depressive-like behaviors (Fig. 1B). Mice treated with ABX did not show significant depressive-like behaviors compared with control mice (Fig. S2A-E). CRS-treated mice showed significant depressive-like behaviors compared to control mice, but these depressive-like behaviors were alleviated after ECS treatment. Interestingly, the ABX treatment of CRS mice abolished the antidepressant effects of ECS, which showed significant increase in immobility in TST and FST, and decrease duration in center of OFT and preference in SPT comparing to CRS mice with ECS treatment (Fig. 1C-G). The ABX treatment of CRS mice did not show significant exacerbation depressive-like behaviors compared to control mice as well (Fig. S2F-J). Collectively, these data suggest that gut microbiota is necessary for the antidepressant effect of ECS in CRS mice.

**Fig. 1.**
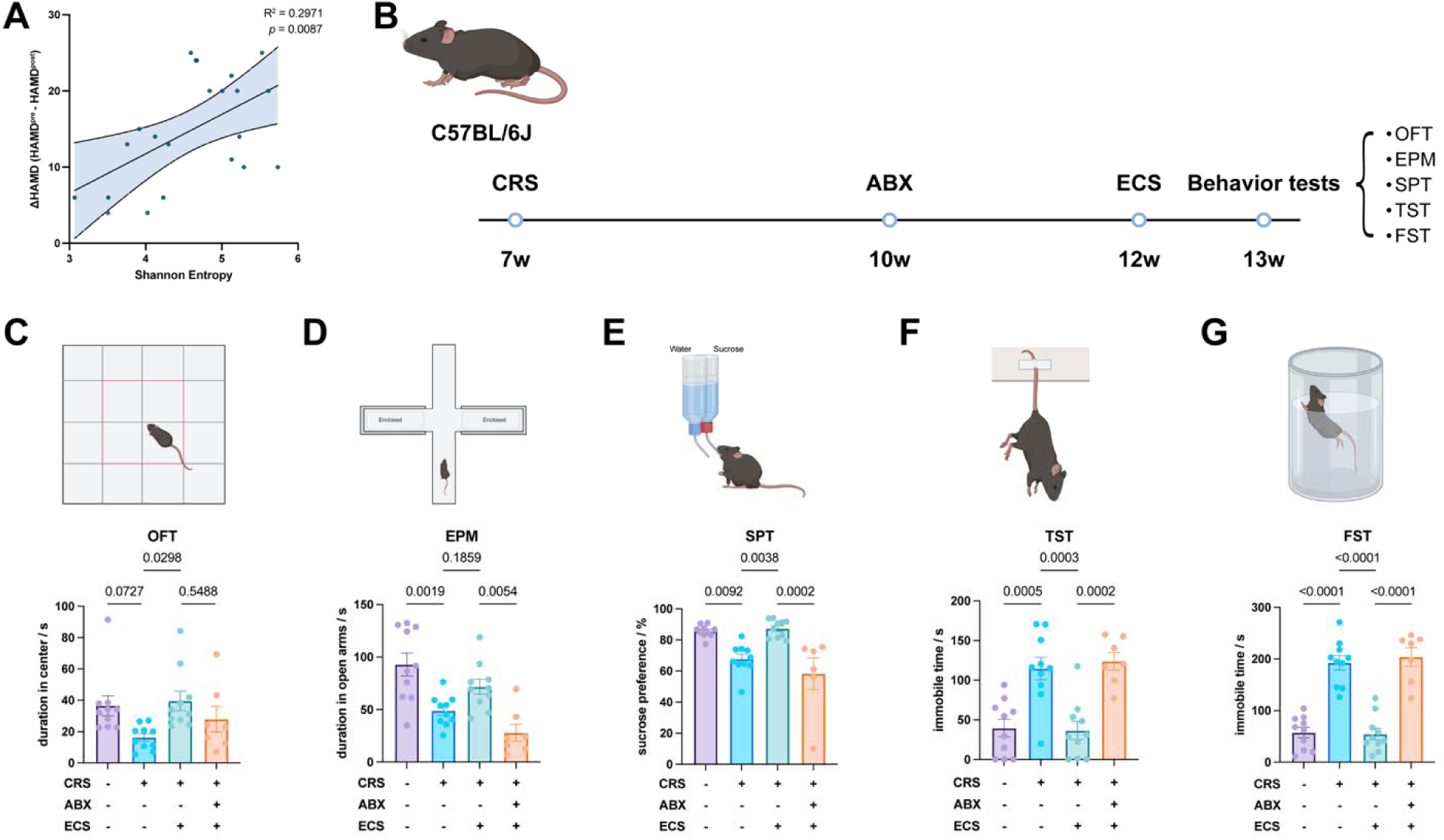
Antibiotics mix (ABX)-induced gut microbiota dysbiosis suppressed antidepressant effect of ECS. A. The decrease in HAMD score after ECT was proportional to the Shannon entropy, an indicator of gut microbiota diversity before ECT in MDD patients, suggesting that gut microbiota dysbiosis in patients with depression may affect the therapeutic effect of ECT. B. Timeline of ABX-induced gut microbiota dysbiosis in the ECS treatment of CRS-induced depression. A CRS model was established in mice aged 7 to 10 weeks. The mice were then treated with ABX or vehicle (VEH) for 2 weeks, followed by 5 days of ECS or sham surgery. At 13 weeks of age, the mice underwent behavioral tests. C. Duration into center area of mice in open-field test (OFT). Mice in CRS-ABX-ECS group showed a decreasing trend in the duration into center area of OFT comparing to mice in the CRS-VEH-ECS group. D. Duration into open arms of mice in elevated plus maze (EPM) test. Mice in CRS-ABX-ECS group showed significantly decreasing duration into open arms of EPM comparing to mice in the CRS-VEH-ECS group. E. The preference for sucrose of mice in sucrose preference test (SPT). Mice in CRS-ABX-ECS group showed significantly decreasing sucrose preference comparing to mice in the CRS-VEH-ECS group. F-G. The immobile time during the test of the mice in the tail suspension test (TST, F) and forced swimming test (FST, G). Mice in CRS-ABX-ECS group showed significantly increasing time of immobility in both TST and FST comparing to mice in the CRS-VEH-ECS group. These results reveal that gut microbiota dysbiosis can significantly reduce the efficacy of ECT treatment for depression. Data are mean ± S.E.M., as determined by ordinary one-way ANOVA with Tukey correction, the significance between groups is expressed numerically.

### Gut microbiota is essential for ECS to regulate the activity of hypothalamus-pituitary-adrenal gland (HPA) axis

The expression of immediate-early genes (IEGs), such as c-Fos, responds to the activation of neurons^[34, 35]^. To screen the neurons that responds to ABX and ECS treatment in CRS mice, a whole brain c-Fos mapping was conducted with a semi-automated procedure. A robust increase of c-Fos expression was observed in paraventricular nucleus of hypothalamus (PVN) of CRS mice compared to control mice. After ECS treatment, the c-Fos expression was recovered. Consistent with previous behavioral results., when CRS mice treated with ABX before ECS, the ECS induced decrease of c-Fos expression was abolished in PVN, and ABX-treated CRS mice didn’t exhibit significant increase of c-Fos expression comparing with ABX-nontreated CRS mice (Fig. 2A-B). The ABX treatment of CRS mice did not show significant activation of neuron in PVN compared to control mice as well (Fig. S2K-L). Further examination of the medial prefrontal cortex (mPFC), nucleus accumbens (NAc), PVN and central amygdala (CeA) of MDD patients were carried out by functional magnetic resonance imaging (fMRI) (Fig. 2C). Based on amplitude of low frequency fluctuation (ALFF)^[36, 37]^ features of the brain regions extracted from the fMRI data, we trained a random forest regression (RFR) model to predict changes of HAMD scores (HAMD^post-ECT^-HAMD^pre-ECT^) of MDD patients. We evaluated the contribution of ALFF features to the RFR model by comparing normalized feature importance of every ALFF feature and found that ALFF features of CeA and PVN contributed most to the model (Fig. 2D). Correlation analysis showed a negative relationship between predicted ΔHAMD (ΔHAMD^predicted^) and actual ΔHAMD of MDD patients, indicating that the model was able to predict the prognosis of ECT for MDD (Fig. 2E).

**Fig. 2.**
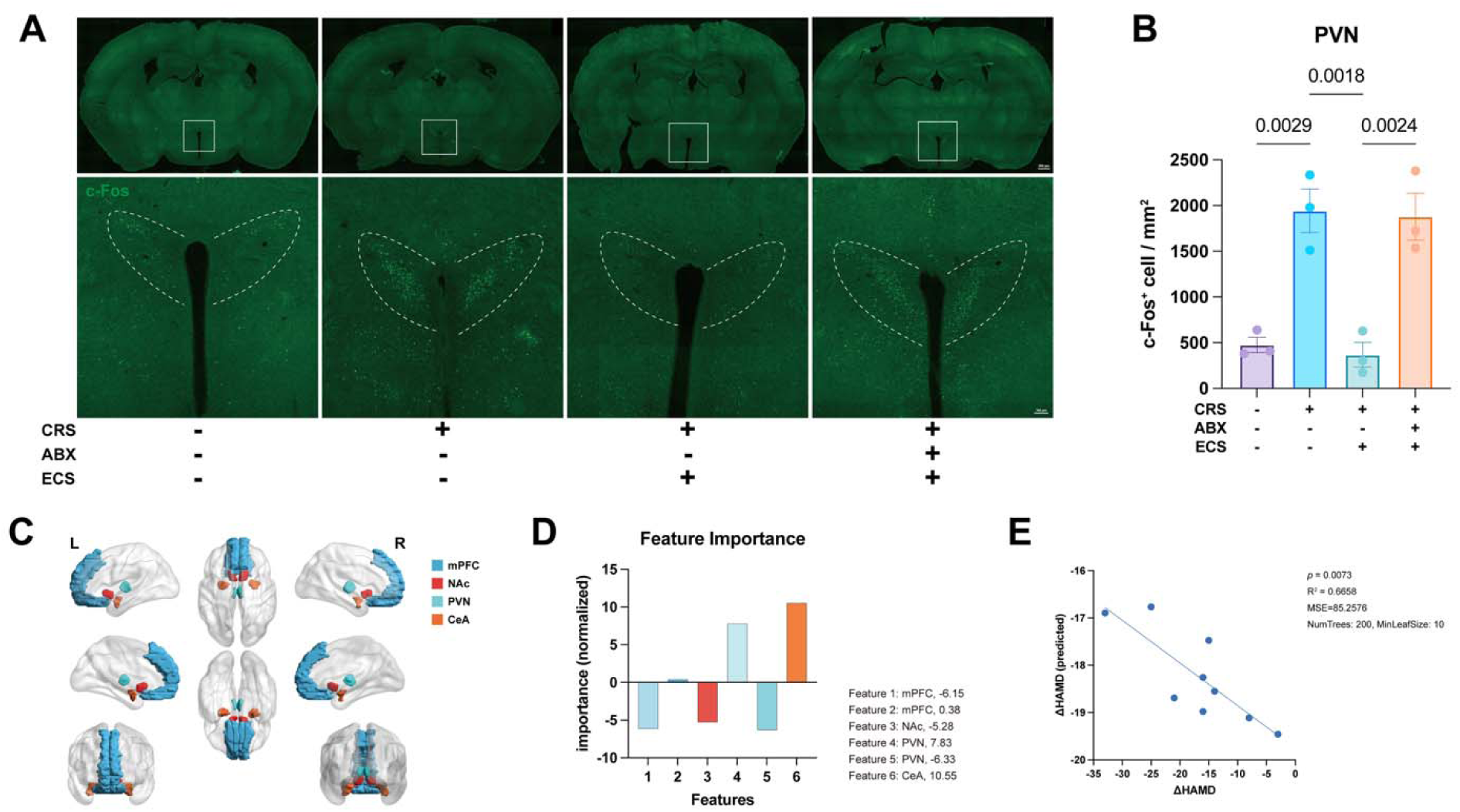
ABX-induced gut microbiota dysbiosis suppressed the downregulation of neuronal activity in paraventricular hypothalamic nucleus (PVN) after ECS. A. Representative imaging of c-Fos, the indicator of neuronal activity, in paraventricular hypothalamic nucleus (PVN) of four groups of mice. B. The numbers of c-Fos-expressing neurons per square-millimeter (mm^2^) in PVN four groups of mice. Compared with the CRS-VEH-SHAM group mice, the density of c-Fos^+^ cells in the PVN of the CRS-VEH-ECS group mice was significantly decreased, however, the density in CRS-ABX-ECS group was remarkably increased comparing to which of mice in CRS-VEH-ECS group. C. Renderings of brain regions of interest in MDD patients included in random forest regression (RFR), the medial prefrontal cortex (mPFC), the nucleus accumbens (NAc), PVN and the central nucleus of the amygdala (CeA) were labeled. D. Normalized importance of ALFF features of brain regions described in RFR model. E. Correlation analysis between predicted changes of HAMD scores (ΔHAMD) and actual ΔHAMD of test database. These results suggest that ECS may exert its antidepressant effects by regulating PVN activity, while ABX treatment may abolish the regulation of PVN activity by ECS. Data are mean ± SEM, as determined by ordinary one-way ANOVA with Tukey correction, the significance between groups is expressed numerically.

Neuroendocrine studies have demonstrated that the over-activated HPA axis leads to elevated cortisol in MDD patients^[38–40]^, and Corticotropin-Releasing Hormone (CRH) neurons in PVN is the driving force of the HPA axis in the central nervous system. Therefore, the serum cortisol of MDD patients were measured before and after ECT to confirm its downregulation of HPA axis. It was observed that a decrease in HAMD scores was positively correlated with a decrease in serum cortisol concentration (Fig. 3A). In addition, the Shannon entropy before ECT was positively correlated with the decrease in serum cortisol concentration, which suggests the effect of ECT in downregulation of HPA axis is negatively correlated with the degree of gut microbiota dysbiosis (Fig. 3B). The significantly increasing number of CRH and c-Fos co-expressing neurons in CRS mice comparing with control mice confirmed overactivation of HPA axis after CRS modeling, and ECS remarkably decrease the number of these co-expressing neurons. Furthermore, ABX treatment abolished the effect of ECS treatment by increasing the number of CRH and c-Fos co-expressing neurons. (Fig. 3C-D). The same result was also observed in serum corticosterone concentration, which was notably increased in CRS mice comparing with control mice, and ECS treatment greatly decreased the serum corticosterone concentration caused by CRS modeling. However, the ABX treatment also abolished the amelioration caused by ECS (Fig. 3E). Taken together, these data suggest that ECS alleviates depressive-like behaviors by downregulating the over-activation of HPA axis, which requires the participation of gut microbiota.

**Fig. 3.**
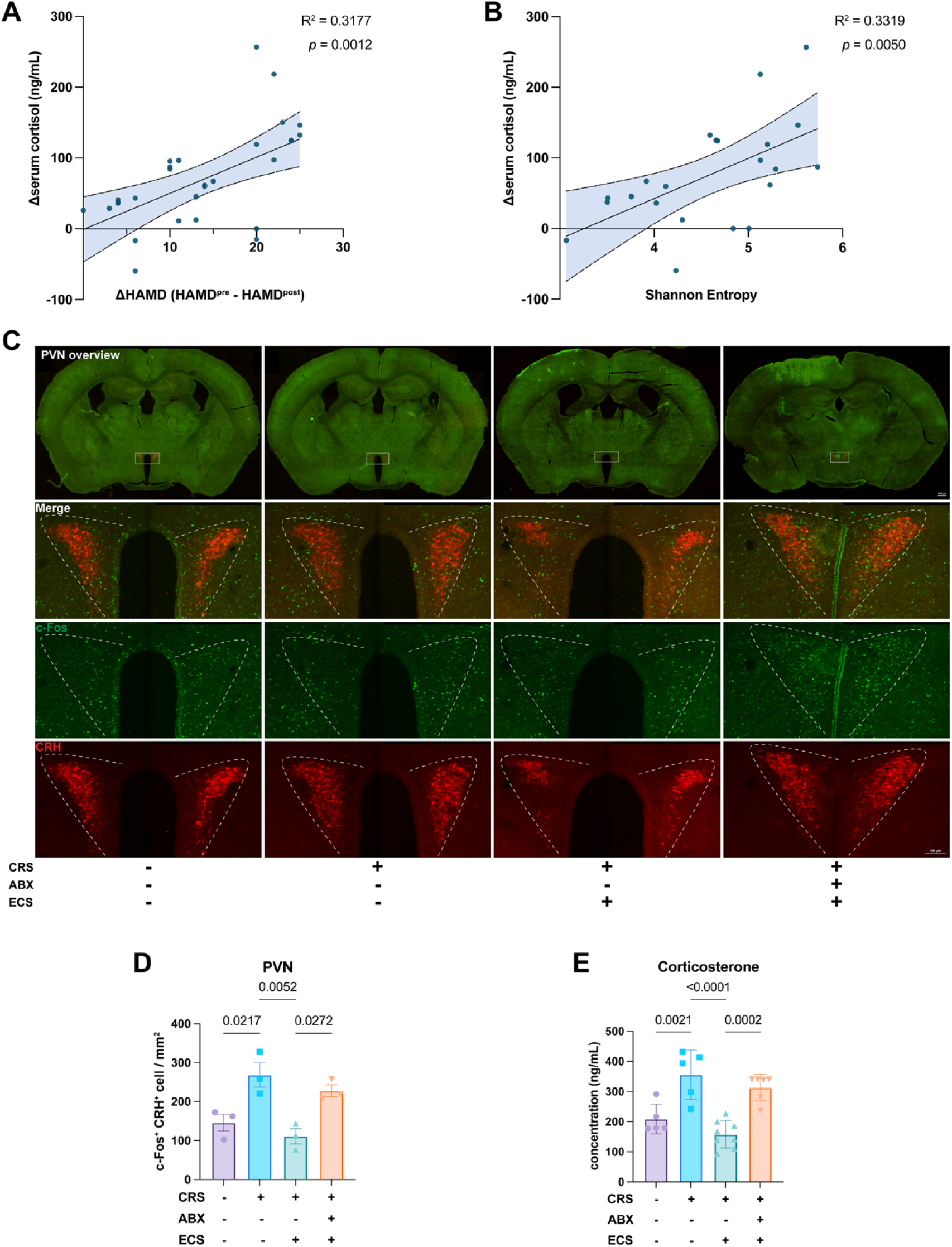
ABX-induced gut microbiota dysbiosis disrupted the regulation of hypothalamus-pituitary-adrenal gland (HPA) axis of ECS. A. The reduction in serum cortisol levels after ECT treatment in MDD patients was positively correlated with the reduction in HAMD scores. B. The reduction in serum cortisol levels after ECT treatment in MDD patients was positively correlated with their Shannon entropy before ECT. C. Representative colocalization imaging of c-Fos and CRH neurons, the activator of HPA axis, in PVN of four groups of mice. D. The numbers of c-Fos-expressing CRH neurons per square-millimeter (mm^2^) in PVN of four groups of mice. Compared with the CRS-VEH-SHAM group mice, the density of c-Fos^+^ and CRH^+^ cells in the PVN of the CRS-VEH-ECS group mice was significantly decreased, however, the density in CRS-ABX-ECS group was significantly increased comparing to which of mice in CRS-VEH-ECS group. E. The serum corticosterone levels of four groups of mice. Compared with the CRS-VEH-SHAM group mice, the corticosterone level of the CRS-VEH-ECS group mice was significantly decreased, however, the corticosterone level in CRS-ABX-ECS group was markedly increased comparing to which of mice in CRS-VEH-ECS group. Thess results confirms that ECS can modulate the HPA axis by regulating CRH neurons in PVN, while ABX treatment may abolish the regulation. Data are mean ± SEM, as determined by ordinary one-way ANOVA with Tukey correction, the significance between groups is expressed numerically.

### Gut microbiota-derived adenosine determines ECS efficacy

To further investigate the mechanisms of gut microbiota in determining ECS efficacy, the metabolites in the feces were compared between the CRS mice treated with ABX or vehicle after ECS by LC-MS. KEGG pathway annotation identified purine metabolism as a potential pathway (Fig. 4A-B), and 4 remarkably downregulated metabolites were observed in the feces of ABX-treated CRS mice after ECS, which were adenosine, adenine, 2’-deoxyadenosine and hypoxanthine (Fig. S3A-D). Notably, adenosine was the most significant metabolite in terms of the magnitude of the decrease, the proportion of change, and the number of metabolic pathways involved in the downregulation (Fig. 4C).

**Fig. 4.**
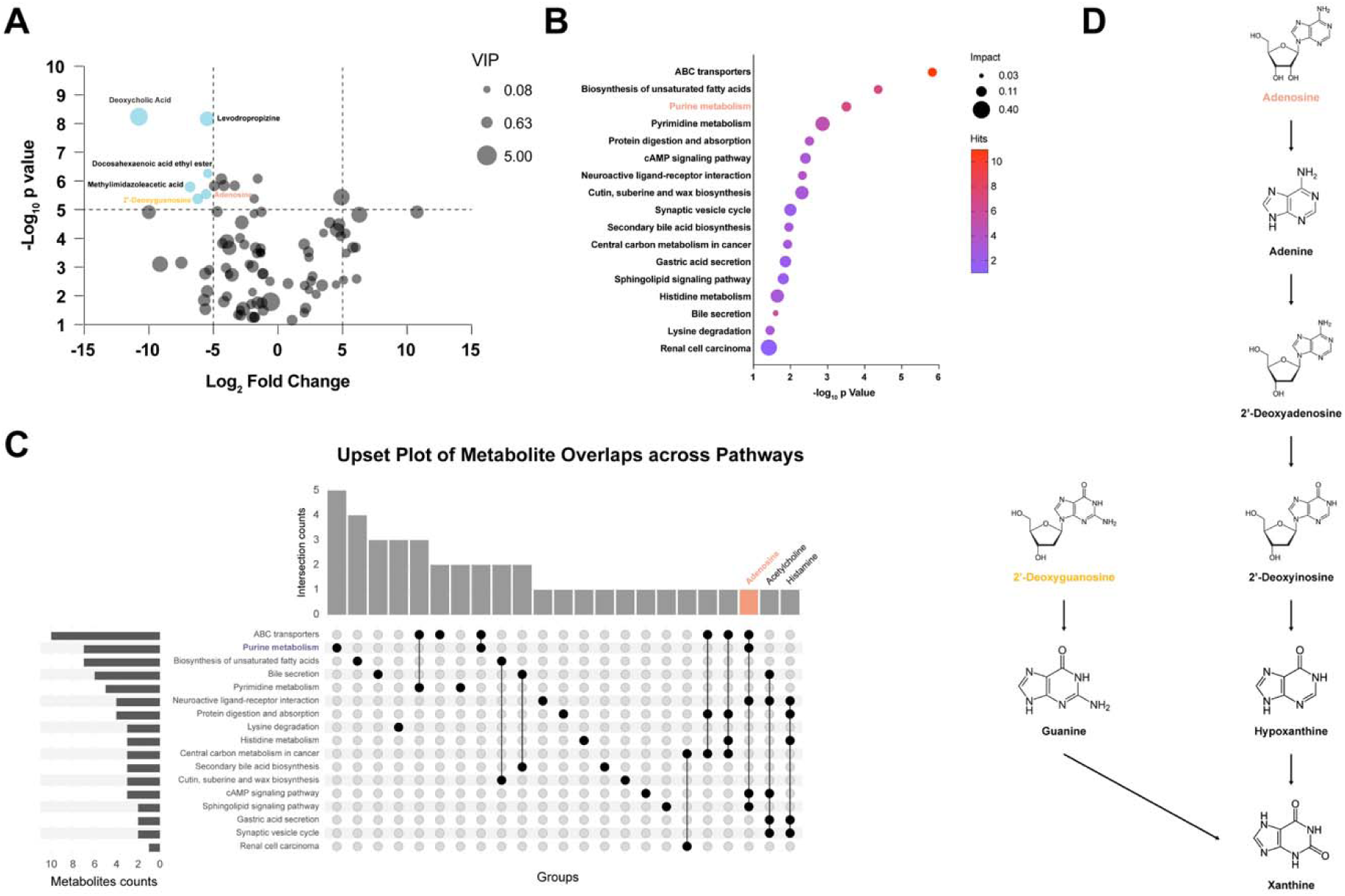
ABX-treatment lowered purine metabolism and adenosine in microbiota of CRS-ECS mice. A. Volcano plot illustrated the distribution of differentially metabolites in the feces between CRS-ABX-ECS and CRS-VEH-ECS mice. The horizontal axis represents the logarithmic fold change in metabolite levels (Log_2_ Fold Change), and the vertical axis represents the adjusted p-value (-Log_10_ p value). The thresholds were set as follows: Log_2_ Fold Change >5 and <-5, and -Log_10_ p value > 5. Above these thresholds, 2’-Deoxyguanosine and Adenosine in the purine metabolism pathway were shown. B. KEGG enrichment analysis of the metabolic pathways of all downregulated metabolites in the feces of CRS-ABX-ECS mice compared to CRS-VEH-ECS mice. The purine metabolism pathway is one of the three pathways that have undergone the most significant changes. C. Upset plot showing the differential metabolites that appeared multiple times in all significantly differential metabolic pathways. Adenosine is one of the three metabolites involved in the most frequent pathways of metabolism. D. A partial schematic diagram of the purine metabolism pathway, showing the relationship of the four metabolites: adenosine, adenine, 2’-deoxyadenosine and hypoxanthine. These results suggest that the purine metabolism pathway and its component adenosine may be the key metabolic pathways and metabolites that regulate ECS effectiveness.

Furthermore, the adenosine levels in feces of MDD patients is the only metabolite that has the significant positive correlation with Shannon entropy (Fig. 5A, Fig. S3E-G), we speculate that gut microbiota-derived adenosine might be the key metabolite for ECS efficacy. To improve this speculation, adenosine was gavaged daily for ABX-treated CRS mice for two weeks and assessed their depressive-like behaviors after 5-day ECS (Fig. 5B). After gavage of adenosine for two weeks, the antidepressant response of ECS was restored in CRS mice treated with ABX, as evidenced by significantly increase duration in the central of the OFT and the open arm of the EPM, as well as a higher sucrose preference of SPT (Fig. 5C-E), and markedly decreased immobility in TST and FST (Fig. 5F-G). Besides, gavage of adenosine alone was not able to ameliorate the depressive-like behaviors of ABX-treated CRS mice (Fig. S4Q-U). The other 3 metabolites were also supplied for ABX-treated CRS mice for two weeks (Fig. S4A), and those mice showed no significant difference in antidepressant effects of ECS (Fig. S4B-P). These results demonstrate that adenosine is the key gut microbiota-derived metabolite for maintaining the efficacy of ECS.

**Fig. 5.**
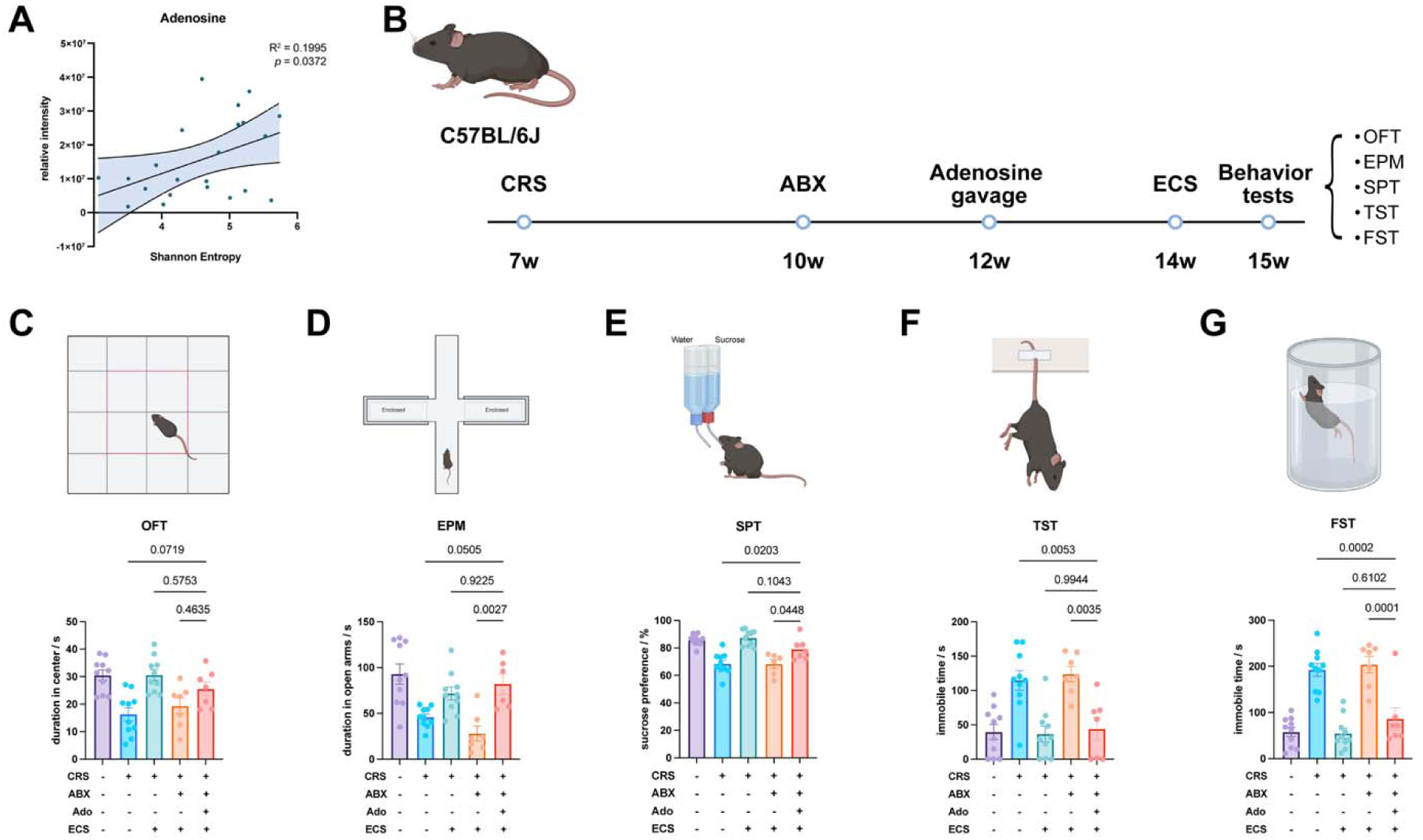
Gut microbiota-derived adenosine sustained efficacy of ECS. A. There was a positive correlation between the relative adenosine level in the feces of MDD patients and their Shannon entropy. B. Timeline of the behavioral tests on metabolite replenishment. A CRS model was established in mice aged 7 to 10 weeks. The mice were then treated with ABX or vehicle for 2 weeks, followed by daily gavage of adenosine (ado) supplements for 2 weeks, and finally underwent ECS or sham surgery for 5 days. Behavioral tests were performed on the mice at 15 weeks of age, and tissue samples were collected after all behavioral tests were completed. C. Duration into center area of mice in OFT. Mice in CRS-ABX-Ado-ECS group showed an increasing trend in the duration into center area of OFT comparing to mice in the CRS-ABX-ECS group. D. Duration into open arms of mice in EPM. Mice in CRS-ABX-Ado-ECS group showed significantly increasing duration into open arms of EPM comparing to mice in the CRS-ABX-ECS group. E. The preference for sucrose of mice in SPT. Mice in CRS-ABX-Ado-ECS group showed significantly increasing sucrose preference comparing to mice in the CRS-ABX-ECS group. F-G. The immobile time during the test of the mice in TST (F) and FST (G). Mice in CRS-ABX-Ado-ECS group showed significantly decreasing time of immobility in both TST and FST comparing to mice in the CRS-ABX-ECS group. These results suggest that adenosine can reverse the decrease in ECT effectiveness caused by ABX-treatment. Data are mean ± S.E.M., as determined by ordinary one-way ANOVA with Tukey correction, n = 7-10, the significance between groups is expressed numerically.

### Adora1-expressing epithelium cell initiated the signaling from gut to the brain

Adenosine signaling is important for regulating depressive-like behaviors^[29, 41, 42]^. It is reported that activation of adenosine A1 receptors (Adora1), which has the highest affinity for adenosine^[43]^, has an antidepressant effect^[27]^. By using transgenic mice, the enteric neuron was labeled by UCHL1-CreERT2 x Ai14 (tdTomato), and Adora1 was marked by immunofluorescence. In the large intestine segment with the most abundant microbiota, confocal imaging showed that enteric nerve fibers could form connections with colon epithelial cells expressing Adora1 (Fig. 6A, S5A). In addition, using herpes simplex virus (HSV) anterograde neural tracing in the colon, fluorescent protein-labeled neuronal cell bodies were observed in the PVN (Fig. 6B). Previous results have shown that ABX treatment can abolish the effects of downregulation on CRS-induced CRH neuron activation and corticosterone concentration decrease by ECS. Notably, supplement with adenosine to CRS-ABX-ECS treated mice could reverse the effects of ABX treatment, which significantly decreased the number of c-Fos and CRH co-expressing neurons in PVN (Fig. 6C-D, S5B-C) and concentration of corticosterone in serum comparing with adenosine non-supplement mice (Fig. 6E).

**Fig. 6.**
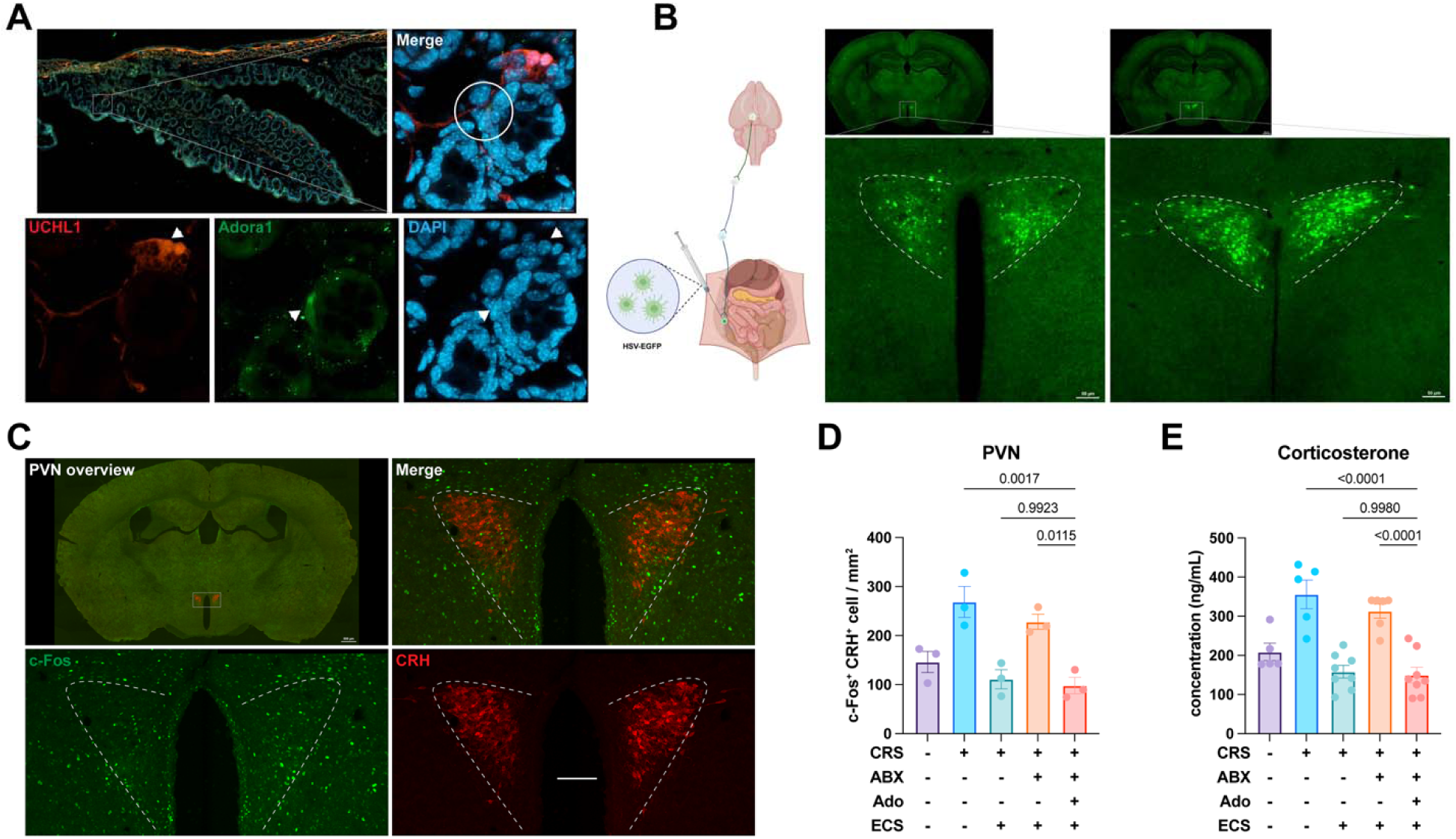
Gut microbiota-derived adenosine may restore the regulatory role of ECS on the HPA axis through the gut-brain axis. A. A fluorescent staining image of intestinal section showing that there may be a connection between adenosine receptor A1(Adora1) expressing-intestinal epithelium cell and UCHL1-labeled enteric neuron. B. Tracing by HSV-EGFP from colon showing that there is a projection from enteric neurons in colon to the neurons in anterior and posterior of PVN. C. Representative colocalization imaging of the c-Fos and CRH neurons in PVN of CRS-ABX-Ado-ECS mice. D. The numbers of c-Fos-expressing CRH neurons per square-millimeter (mm^2^) in PVN of 5 groups of mice. Compared with the CRS-ABX-ECS group mice, the density of c-Fos^+^ and CRH^+^ cells in the PVN of the CRS-ABX-Ado-ECS group mice was significantly decreased, which has no difference in density compared with the CRS-VEH-ECS group mice. E. Serum corticosterone concentration of 5 groups of mice. Compared with the CRS-ABX-ECS group mice, the corticosterone level of the CRS-ABX-Ado-ECS group mice was markedly decreased, which has no difference in corticosterone level compared with the CRS-VEH-ECS group mice. These results imply that gut microbiota-derived adenosine may act on the CRH neurons in PVN to regulate HPA axis by gut-brain axis. Data are mean ± SEM, as determined by ordinary one-way ANOVA with Tukey correction, the significance between groups is expressed numerically.

According to these results, we surmise that adenosine secreted by gut microbiota may acts on intestinal epithelial cells through the Adora1 receptor, then transmit the signal to PVN in the central nervous system through the enteric neurons, forming a microbiota-epithelial-neural pathway to regulate the activation of HPA axis, which may determine the efficacy of ECS.

### Gut microbiota-derived adenosine limits the adverse effects of ECS

Retrograde amnesia is the most common persistent adverse effect of ECT^[1, 12]^. The clinical data showed a significant decrease of Mini-Mental State Exam (MMSE)^[44]^ scores, a measurement of cognitive level, in MDD patients with gastrointestinal symptoms after ECT (Fig. 7A), which suggested gut microbiota may influence the adverse effect of ECT on memory. Considering depression affects cognitive function, healthy mice were uses instead of CRS mice for experiments of following adverse effect study. After adapted for 1 week, mice were treated ABX or vehicle for 2 weeks, and their cognitive functions were measured by Morris water maze (MWM) test after 5-day ECS treatment (Fig. S6A). After ECS, mice exhibited impaired spatial memory, as evidenced by a significant increase in escape latency and decrease of staying time in target area by comparing to non-ECS treated mice in MWM test. ABX treatment alone didn’t impair spatial memory. However, when ABX was treated before ECS, it aggravated the impairment of ECS on spatial memory (Fig. S6B-C). To investigate the changes of synaptic plasticity in the central nervous system induced by ABX and ECS treatment, which is one of the key indicators of learning and memory ability, the density of PSD95 was examined in medial prefrontal cortex (mPFC), Dentate gyrus (DG) and Caudoputamen (CPu) (Fig. 7E). ECS led to remarkably reduced density of PSD95 in mPFC and DG, which are highly correlated with memory formation, but no significant difference was observed in CPu, which is correlated with motor function. Additionally, the density of PSD95 in DG of mice treated with ABX and ECS was even lower than the mice treated with ECS along (Fig. 7F-H). Overall, these data indicated that gut microbiota may play a protective role against memory impairment during ECT treatment.

**Fig. 7.**
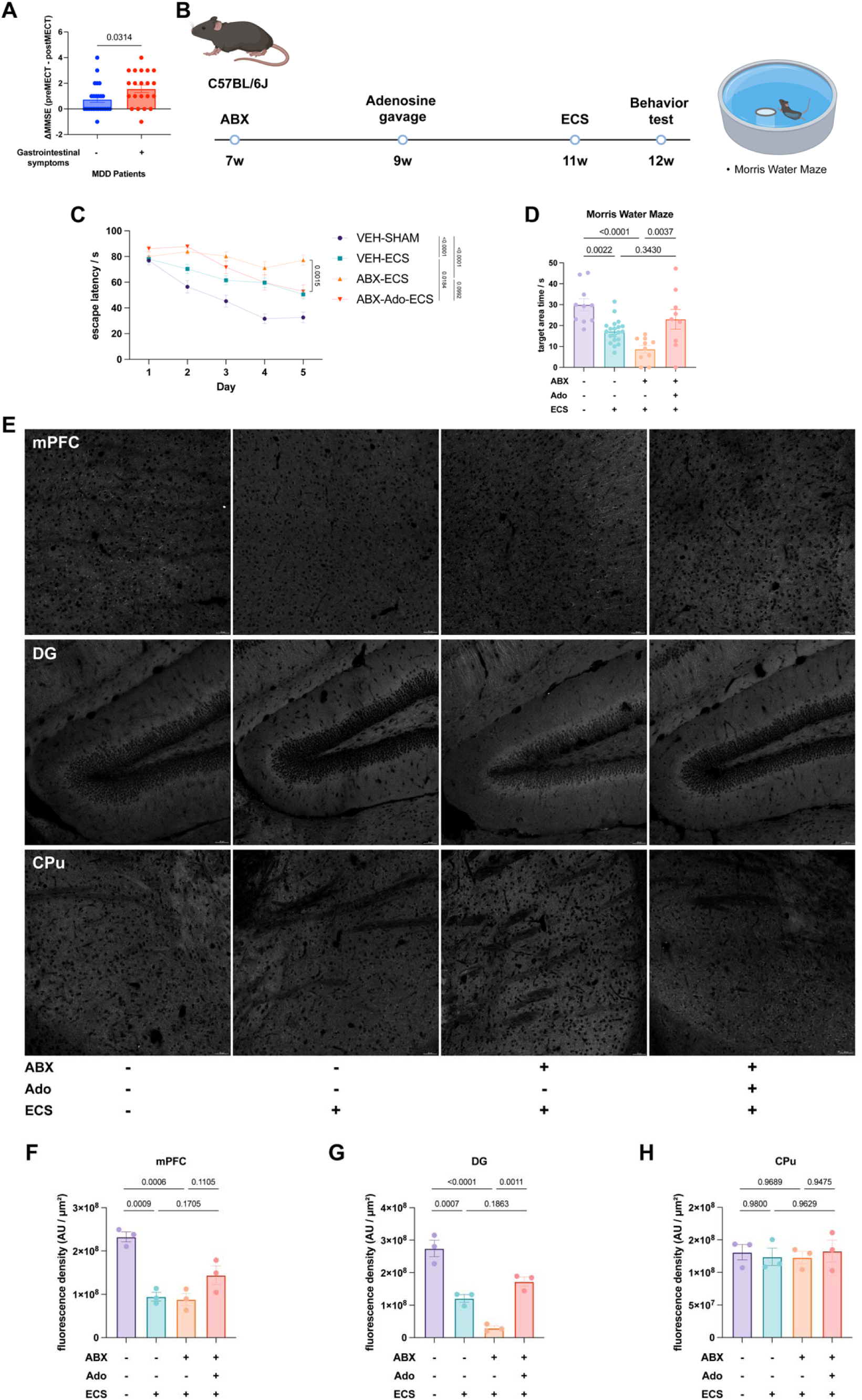
ABX-induced microbiota dysbiosis exacerbated memory loss after ECS, which could be ameliorated by adenosine supplementation. A. MDD patients with gastrointestinal symptoms showed a significantly greater decline in the MMSE cognitive score after ECT than patients without gastrointestinal symptoms. B. Timeline of experiments on metabolite replenishment to alleviate memory loss. The mice were treated with ABX or vehicle for 2 weeks, followed by daily gavage of adenosine supplements for 2 weeks, and finally underwent ECS or sham surgery for 5 days. Behavioral tests were performed on the mice at 12 weeks of age. C. Escape latency in acquisition period of all groups of mice by day. Compared with the mice in VEH-SHAM group, mice in both the VEH-ECS and ABX-ECS groups showed dramatically prolonged escape latency. By supplying adenosine, mice in the ABX-Ado-ECS group showed a significant decrease in escape latency on day 5 compared to mice in the ABX-ECS group. D. Time lingering in target area of all groups of mice at the last day of testing. By supplying adenosine, mice in the ABX-Ado-ECS group showed a significant increase in duration of stay in the target area compared to mice in the ABX-ECS group. E. Representative imaging of PSD-95 of medial prefrontal cortex (mPFC), hippocampal dentate gyrus (DG) and caudoputamen (CPu) of ABX-Adenosine-ECS mice. F-H. Density of fluorescence intensity of PSD-95 of mPFC (F), DG (G) and CPu (H) of all groups of mice. By comparing the PSD95 density in the mPFC, DG, and CPu of mice in ABX-ado-ECS group and ABX-ECS group, adenosine supplementation significantly increased the PSD95 density in DG, and showed an increasing trend in mPFC, but had no effect in CPu. These results indicate that adenosine supplementation can alleviate the further deterioration of post-ECS memory loss induced by ABX-treatment. Data are mean ± SEM, as determined by ordinary one-way ANOVA with Tukey correction, the significance between groups is expressed numerically.

In order to explore whether intestinal metabolites play a protective role on the memory impairment induced by ECS, the metabolites were supplied to the ABX-treated mice for 2 weeks and MWM test was performed after 5-day ECS (Fig. 7B, S6D), the supplement of adenosine and 2’-deoxyadenosine to ABX-treated mice limited spatial memory impairment after ECS compared to ABX-ECS group (Fig. 7C-D, S6E H), while the supplement of adenine and hypoxanthine showed no significant results (Fig. S6F-G I-J). Meanwhile, the density of PSD95 was also examined in mPFC, DG and CPu with adenosine replenishment (Fig. 7E). An upregulation of PSD95 density in the DG and mPFC were observed comparing to the density of the ABX-ECS treated. (Fig. 7F-H). Overall, these data indicated that gut microbiota and microbiota-derived adenosine may play a protective role against memory impairment during ECT treatment.

## Discussion

Gut microbiota has been shown to influence MDD and other affective disorders^[45–48]^. Evidence from observational studies and meta-analyses suggests a significant role of gut microbiota in mediating mental health via probiotic administration and affect efficacy of drugs or other medication^[46, 49]^. A study observed that administration of a probiotic formulation consisting of *Lactobacillus helveticus* R0052 and *Bifidobacterium longum* R0175 significantly reduced anxiety-like behaviour in rats and alleviated psychological distress in volunteers^[50]^; and many other clinical researches reported antidepressant efficacy of probiotics, including *Lactobacillus*, *Bifidobacterium* and their formulations^[46]^. Studies on duloxetine, a widely used antidepressant, was found to be bioaccumulated by various pathogenic strains such as S*treptococcus salivarius*, *B. uniformis* and *E. coli IAI1* but not bio-transformed, which might impair the efficacy of duloxetine by decreasing its bioavailability^[47]^. As a procedure which can greatly and rapidly improve MDD symptoms and other mental disorders, some case reports have indicated connections between microbiota and ECT as well^[19, 21]^, for example, clinical study on patients with severe or treatment-resistant depression has reported a significant difference of alpha diversity of oral microbiota between ECT responders and non-responders^[21]^. Yet, mechanisms of ECT and its interaction with gut microbiota are still elusive. Our study observed significant impairment of antidepressant effect of ECS in ABX-treated CRS mice, suggesting gut microbiota and its metabolite adenosine determines ECT efficacy.

The HPA axis is a critical endocrine system that orchestrates the stress response, which involves a complex set of behavioral, neuroendocrine, autonomic, and immune responses to enable adaptation to aversive psychological and physiological stimuli^[39, 51]^. The main components of the HPA axis are PVN, the anterior lobe of the pituitary gland, and the adrenal cortex^[39]^. In the HPA axis, PVN releases CRH to stimulate the anterior pituitary to release adrenocorticotropic hormone (ACTH) into the peripheral circulation, which stimulates the adrenal glands to release glucocorticoids (e.g., cortisol, or corticosterone). Glucocorticoids act on both glucocorticoid receptors and mineralocorticoid receptors, and regulate other parts of the HPA axis and higher brain structures (e.g., cortex, hippocampus, and periventricular thalamus) by negative feedback^[39, 51]^.It is reported that patients with MDD exhibited hyperactivity of the HPA axis with impaired negative feedback^[40, 52]^. We found that ECS remarkably decrease the numbers of CRH and c-Fos co-expressing neuron and the serum corticosterone levels in CRS mice, and that ABX treatment abolished these effects of ECS treatment. These results suggest that gut microbiota is required for ECT to alleviates depressive-like behaviors by reducing the activation level of the HPA axis.

Adenosine is a naturally substance that relaxes and dilates blood vessels, and affects the electrical activity of the heart as well^[53]^. All adenosine-containing substances, such as adenosine phosphate, S-adenosine methionine, S-adenosine homocysteine, and RNA, are capable of generating adenosine during their degradation; but the most primarily way to generate adenosine is dephosphorylating adenosine phosphate. Recent studies have identified a metabolic pathway for the microbial origin of adenosine^[54]^, and our observation of decreased adenosine level in feces of mice after ABX-induced dysbiosis supports this finding. Adenosine has been used for its antiarrhythmic properties in supraventricular tachycardia (SVT), and it can also function as a diagnostic tool, depending on the type of SVT^[53]^. In addition, adenosine is a prominent physiological mediator of sleep homeostasis, which is released in the basal forebrain (BF), a brain region that plays a critical role in regulating the sleep-wake cycle, to suppress neural activity mediated by the A1 receptor and increase sleep pressure^[55, 56]^. A metabolic profiling of plasma to explore the potential biomarkers of depression in children and adolescents with MDD has identified several perturbed pathways, including fatty acid metabolism and purine metabolism, that are associated with MDD in these young patients, and adenosine is decreased in plasma from children and adolescents with MDD^[57]^. Other experimental studies also suggest that adenosine is critical for alleviating depressive-like behaviors^[29, 40]^. We found that supplementation with adenosine could reverse the effects of ABX treatment, which recovered antidepressant effects of ECS, and significantly decreased the number of c-Fos and CRH co-expressing neurons in PVN and concentration of corticosterone in serum comparing with CRS-ABX-ECS treated mice. These results suggest that gut microbiota-derived adenosine may be the key metabolite that determines ECS efficacy by regulating HPA axis activity.

## Methods

### Mice

All animals (C57BL/6J (B6), male, aged 6–7 weeks) used in this study were purchased from Gempharmatech, China. Unless otherwise noted, all the mice were housed in groups of four/five and kept on a 12/12 h light/dark cycle in a temperature-controlled environment (22 ±1°C) and had free access to food and drinking water throughout the experiment. All experiments were conducted during the light cycle. All mice were maintained under specific pathogen-free (SPF) conditions and kept at a strict 24 h light-dark cycle, with lights on from 8 a.m. to 8 p.m. in the animal facility of the University of Science and Technology of China. All animal experiments were approved by the Ethics Committee of University of Science and Technology of China (USTCACUC28120125096). All efforts were made to minimize animal suffering as well as the number of animals used.

### Patient Inclusion and Exclusion Criteria

A total of 111 patients with major depressive disorder were recruited from Anhui Mental Health Center between June 2013 and March 2022. Diagnosis was conducted by two psychiatrists based on the Diagnostic and Statistical Manual of Mental Disorders-5 (DSM-5) criteria. Exclusion criteria were predefined as follows: (1) age younger than 18 years or older than 65 years; (2) pregnancy, substance abuse, severe physical illness, current or past neurological disorders, or other psychiatric comorbidities (e.g. schizophrenia, bipolar I or II disorder, and Axis II disorders) assessed using the Structured Clinical Interview for DSM-5 (SCID-5)^[58]^; (3) received fewer than 5 or more than 12 ECT sessions (n=6). Ultimately, 111 participants were included in this study.

This study was conducted in accordance with the principles of the Declaration of Helsinki. Ethical approval was obtained from the local ethics committee of Anhui Medical University (Study No. 84230095). Written informed consent was obtained from all participants before their inclusion in the study.

### Electroconvulsive Therapy (ECT) Procedure

ECT was collaboratively administered by experienced psychiatrists and anesthesiologists using the Thymatron System IV device at the Anhui Provincial Mental Health Center. Prior to treatment, patients underwent comprehensive preoperative evaluations to confirm the absence of contraindications for anesthesia or ECT. The treatment team provided patients and their families with comprehensive information concerning the potential benefits and risks, and obtained informed consent. Patients were instructed to discontinue anticonvulsant medications (e. g. benzodiazepines) and to fast before the procedure.

During the procedure, patients received anesthesia under continuous electrocardiogram (ECG) monitoring and adequate respiratory support by the anesthesia team. The induction of anesthesia was carried out using propofol at a dose of 1.4 mg/kg. Muscle relaxation was achieved by administering succinylcholine at 0.5 mg/kg, and atropine (0.5 mg) was given to reduce glandular secretions. After anesthesia, bilateral electrodes were placed on the frontal regions, delivering a 1 ms short pulse at 90 mA for approximately 5 seconds, with energy adjusted based on patients’ age. Closely observation was performed until patients regained consciousness and vital signs were stable after ECT. The initial three treatments were administered once a day; the following treatments were given every other day. The total number of sessions was determined by the treatment team based on patients’ response, typically ranging from 6 to 12 sessions. Patients’ psychiatric medications remained unchanged throughout the entire sessions.

### Clinical assessment and Tissue collection

The MDD patients were interviewed face-to-face by trained psychiatrists. To improve the objectivity and reliability of the measurements, all the interviewers were psychiatrists with at least 5 years of experience in clinical practice and were well-trained for this project. Basic socio-demographic and clinical data were collected, and participants completed a self-administered questionnaire. The questionnaire was administered at baseline, 12-24 hours before the first ECT session, and at follow-up, 24–72 hours after the final ECT session. Human blood and fecal samples were collected from participants in accordance with ethical guidelines.

Depression severity was evaluated using the Hamilton Depression Rating Scale (HAMD-17)^[30]^. A custom-designed questionnaire was used to assess gastrointestinal symptoms and constipation-related issues, with patients completing it under the guidance of trained psychiatrists. Cognitive function was assessed using the Mini-Mental State Examination (MMSE), a widely used screening tool for cognitive impairment^[44]^. The MMSE consists of 30 items evaluating multiple cognitive domains, including orientation, attention, memory, language, and visuospatial abilities. Higher scores indicate better cognitive performance, with a maximum possible score of 30. Patients completed the MMSE under the guidance of trained psychiatrists. Patients completed the Gastrointestinal Symptom Rating Scale (GSRS), a self-administered questionnaire consisting of 15 multiple-choice items^[31, 59]^.

The GSRS assesses five gastrointestinal symptom domains: abdominal pain (including abdominal pain, hunger pains, and nausea), reflux syndrome (heartburn and acid regurgitation), diarrhea syndrome (diarrhea, loose stools, and an urgent need for defecation), indigestion syndrome (borborygmus, abdominal distension, eructation, and increased flatus), and constipation syndrome (constipation, hard stools, and a feeling of incomplete evacuation)^[60]^.

### MRI Acquisition

All participants underwent MRI scans in the dedicated MRI suite of the Medical Imaging Center at the University of Science and Technology of China, using a 3.0T GE 750 MRI scanner (Discovery GE750w, GE Healthcare, Buckinghamshire, UK). Prior to scanning, participants were thoroughly briefed on the procedure and safety precautions, and screening for MRI contraindications, such as metal implants, was conducted. To minimize noise exposure, participants wore earplugs and had foam padding placed around their heads to reduce motion. They were instructed to maintain a comfortable position, remain still, keep their eyes closed, relax, and stay awake.

Scanning Protocols : high-resolution T1-weighted images were acquired using a 3D-BRAVO sequence with the following parameters: repetition time (TR) = 8.16 ms, echo time (TE) = 3.18 ms, inversion time (TI) = 450 ms, flip angle = 12°, matrix size = 256×256, field of view (FOV) = 256 mm × 256 mm, voxel size = 1×1×1 mm³, slice thickness = 1 mm, 188 slices. Resting-state functional MRI (rs-fMRI) was performed using an echo-planar imaging (EPI) sequence with TR = 2400 ms, TE = 30 ms, flip angle = 90°, matrix size = 64×64, FOV = 192 mm × 192 mm, voxel size = 3×3×3 mm³, slice thickness = 3 mm, 46 slices per volume, totaling 9982 volumes.

This study collected imaging data from 45 major depressive disorder (MDD) patients before and after ECT treatment.

### Magnetic Resonance Data Processing Resting-State fMRI Data Preprocessing

Resting-state fMRI data were preprocessed using SPM12 and DPABI 6.0 on the MATLAB platform. The steps included: (1) discarding the first 10 time points to reduce initial magnetic instability; (2) slice timing correction; (3) head motion correction, excluding subjects with displacement >3 mm/3° or mean FD >0.5 mm; (4) spatial normalization to MNI space with resampling to 3 × 3 × 3 mm³; (5) smoothing with a 4 mm FWHM Gaussian kernel; (6) linear trend removal; (7) regressing out Friston-24 head motion parameters, time points with FD >0.5 mm, and signals from cerebrospinal fluid, white matter, and whole brain; (8) temporal filtering to retain 0.01-0.1 Hz signals.

Feature data, such as ALFF, were extracted from the prefrontal cortex, the hypothalamic paraventricular nucleus, nucleus accumbens, and central amygdala. The difference in ALFF values before and after ECT formed the feature set. Missing values in rating scales were imputed with the mean, and all features were standardized using Z-score normalization.

### Random Forest Regression, RFR

RFR was used to predict HAMD scores based on fMRI-derived brain region features. Model performance was evaluated by the correlation coefficient between predicted and actual HAMD scores. Feature importance for each ALFF indicator was standardized, with total importance normalized to 1 for direct comparison.

Compared to multiple Pearson correlations, which require strict correction for multiple comparisons and may increase false negatives, machine learning models are better suited to high-dimensional data and can identify key brain regions influencing the outcome. RFR, as an ensemble method, improves robustness and accuracy by averaging multiple regression trees, making it ideal for this study’s multi-feature data^[61]^.

Model hyperparameters were optimized using grid search and five-fold cross-validation, with final settings: 300 trees, minimum leaf size of 10, and maximum features set to the square root of the total feature number. Data were split 80% for training and 20% for testing. Evaluation metrics included mean squared error (MSE), R², correlation coefficient (r), and p-value (p < 0.05). Analyses were performed using Python’s scikit-learn, with data processing and visualization via Pandas and Matplotlib.

### Chronic Restraint Stress (CRS)

Mice were individually restrained in a well-ventilated 50-ml conical tube for 8h daily from 9 a.m. to 5 p.m., and this procedure was repeated for 21 days. After each daily restraint session, the mice were placed in their home cages with free access to food and water^[62]^.

### ABX treatment

Mice were supplied with a mixture of antibiotics (ampicillin (Amp) 1mg/mL, neomycin (Neo) 1mg/mL, metronidazole (Met) 1mg/mL and vancomycin (Van) 0.5mg/mL) in drinking water for 14 days. Fresh feces samples were collected for the measurement of fecal bacteria load^[63]^.

### Metabolite treatment

Mice were given adenosine (Ado, 50 mg/kg body weight), adenine (A, 25 mg/kg body weight), 2’-deoxyadenosine (dAd, 12.5 mg/kg body weight), or hypoxanthine (I, 100 mg/kg body weight) individually daily for 14 days by gavage.

### Electroconvulsive Shock

ECS was administered using auricular electrodes (Ugo Basile SRL, Gemonio VA, Italy) under anesthesia^[64]^. Mice were treated with 2% isoflurane. Thirty minutes later, ECS (100 Hz, 12 mA, pulse width 0.5 ms, duration 1 s) was administered using an electrical stimulator and an isolator. ECS induced visible tonic clonic seizures lasting for 10-20 s. ECS was performed once daily, 5 times in 5 days. For mice in the sham groups, only 2% isoflurane was administered using similar auricular electrodes without electrical stimulation^[65]^.

### Open Field Test (OFT)

The open field test was conducted in a rectangular plastic box (50 cm × 50 cm × 50 cm) divided into 16 (4 × 4) identical sectors (12.5 cm × 12.5 cm). The field was subdivided into peripheral and central sectors, where the central sector included 4 central squares (2 x 2), and the peripheral sector was the remaining squares. The mouse was placed into the center of an open field and allowed to explore for 6 min. The apparatus was thoroughly cleaned with diluted 75% ethanol between each trial. A video tracking system (Shanghai Xinruan Information Technology Co., Ltd; China) was used to analyze locomotor activity by tracking the distance traveled. Time spent and entries into the center were measured as indicators of anxiolytic behavior^[66]^.

### Elevated Plus Maze (EPM)

The Elevated Plus Maze (EPM) test was used to assess both anxiety-like and depressive-like behaviors in mice. The maze consisted of two open arms without walls and two enclosed by 15 cm high walls, and each arm was 30 cm long and 6 cm wide. Each arm of the maze was attached to sturdy legs such that it was elevated 50 cm of the ground. Mice were placed at the center facing an open arm and allowed to explore freely for 6 minutes. The percentage of time spent in the open arms and the number of open arm entries were recorded, with both reduced open-arm exploration and fewer open-arm entries indicating increased anxiety- and depressive-like behavior.

### Sucrose Preference Test

Mice were acclimated to two bottles of water for 24 hours while single-housed, followed by a 24-hour water deprivation period. Subsequently, a 48-hour testing period (starting at 14:00) involved providing mice with one bottle of 1% sucrose (w/v) and one bottle of water. The position of the two bottles was switched at 24h. The percentage of sucrose preference was calculated as the following calculation method: Sucrose preference% =[(the consumption of 1% sucrose/the total liquid consumed (water and sucrose)] x 100^[67]^.

### Tail Suspension Test

The mouse was suspended by their tails from a lever in a 30 cm × 15 cm × 15 cm opaque white chamber. Animal behavior was recorded using a side camera right after the mouse was suspended for 5 minutes in a quiet environment^[68]^, and immobility duration was quantified by two blinded, trained experimenters. Data were presented as immobility time across different groups.

### Forced Swim Test

Mice were placed in a clear plastic cylinder-shaped bucket (30 cm tall, 20 cm diameter) filled with water at a height that their tails were not able to reach the bottom of the bucket at 30 ± 1 °C. Mice were allowed to swim for 6 min, and their behaviors were recorded by a video camera placed in front of the tank allowing the clear observation of the animals’ behavior later on while viewing the footage. Immobility was quantified independently by two blinded, trained experimenters. Code the duration spent as “Immobile” if the mouse is floating without any movement except for those necessary for keeping the nose above water. Code the duration of time spent as “Swimming” if the movement of forelimbs or hind limbs in a paddling fashion is observed. Data were expressed as immobility time (min) in different groups^[69]^

### Morris Water Maze (MWM) Test

The MWM is a circular tank (120 cm in diameter and 50 cm in height) with non-reflective interior surfaces. The circular pool was filled with water opaqued with a nontoxic water-soluble white dye (titanium dioxide) at 20LJ±LJ2LJ°C and was divided into 4 quadrants of equal area. A platform (9 cm in diameter and 15 cm in height) submerged 1 cm below the surface of water was centered in one of the 4 quadrants of the pool so that it was invisible at water level, and the position of the platform remained constant. During the training session, mice were subjected to swim training for 90 s to find the platform. The mice were given 4 trial sessions each day for 5 consecutive days, with an inter-trial interval of 15 min, and the escape latencies were recorded. This parameter was averaged for each session of trials and for each mouse. Once the mouse located the platform, it was permitted to remain on it for 30 s. If the mouse was unable to locate the platform within 90 s, it was placed on the platform for 30 s and then returned to its cage by the experimenter. On day 6 (test session), the probe trial test involved removing the platform from the pool and mice were allowed the cut-off time of 120 s. Animal behaviors were all recorded by a video camera above the MWM^[70]^.

### Metabolite extraction from fecal samples

Metabolites were extracted from less than 30 mg of sample in 50 volumes per weight solvent (methanol: acetonitrile: MilliQ water = 2: 2: 1 (v/v/v)). Samples were vortexed for 30 s, homogenized for 2 min in a Servicebio tissue homogenizer at maximal speed (60Hz), undergone a 10-min ultrasonication 3 times and incubated at -20℃ overnight. On the next day, samples were centrifuged for 15 min at 13,000 rpm at 4℃. From the supernatant, 200μL were transferred to 1.5 mL centrifuge tubes and undergone vacuum evaporation and concentration using a CV600 vacuum centrifuge concentrator (Beijing Jiaimu Technology). and stored at -20℃. For mass spectrometry analysis, the samples were diluted 1:1 in mobile phase (phase A (acetonitrile: MilliQ water: 1M ammonium formate = 50: 49: 1 (v/v/v), 0.1% formic acid (v/v)): phase B (acetonitrile: MilliQ water: 1M ammonium formate = 95: 4: 1 (v/v/v), 0.1% formic acid (v/v)) = 1: 1 (v/v)), and stored at -20℃ until analysis.

### LC–MS-based untargeted and targeted metabolomics

Samples were analyzed using an Orbitrap Exploris 120 mass spectrometer coupled with a Vanquish Flex UHPLC system (Thermo Fisher Scientific). Chromatographic separation was achieved on an ACQUITY Premier BEH Amide column (2.1 × 150 mm, 1.7 μm; Waters, Cat# 186009506) with a 2 μL injection volume. A linear gradient elution was performed at 0.3 mL/min using mobile phase A (0.01% formic acid [FA], 10 mM ammonium formate [NH4FA] in ACN:H2O 5:95, v/v) and B (0.01% FA, 10 mM NH4FA in ACN:H2O 95:5, v/v), with phase B increasing linearly from 2% to 98% over 16.0 min. The system was re-equilibrated with 2% B for 2.0 min between runs.

Mass spectrometry data were acquired in positive or negative ionization modes, covering a mass range of m/z 70-1050 in both full-scan and data-dependent MS² (ddMS²) acquisition modes. Electrospray ionization parameters were optimized as follows: ion spray voltage, 3.4 kV (positive mode) and 2.3 kV (negative mode); sheath gas flow rate, 40 arbitrary units (arb); auxiliary gas flow rate, 15 arb; ion transfer tube temperature, 320°C; vaporizer temperature, 300°C.

The acquired raw data were processed using Compound Discoverer 3.3 (Thermo Fisher Scientific) and TraceFinder 5.1 (Thermo Fisher Scientific) software packages. Qualitative identification was performed by matching observed molecular ion peaks (m/z) with authentic certified reference standards, while quantification was conducted using validated multi-point calibration curves of target analytes (adenosine [m/z 268.104], adenine [m/z 136.062], hypoxanthine [m/z 137.046], and deoxyadenosine [m/z 252.109]). All samples were analyzed in triplicate to verify method stability and reproducibility, with quantification performed via external standard calibration. Rigorous quality control measures included systematic instrument cleaning procedures and regular blank analyses interspersed between sample runs to eliminate potential cross-contamination and solvent carryover effects, thereby ensuring both data integrity and instrumental reliability throughout the analytical sequence.

### Corticosterone Measurement

Mice were sacrificed and blood was collected between 09:00 a.m. and 11:00 a.m. 2 days after the behavior tests. The members of different groups were collected randomly. Blood samples were allowed to coagulate for 30 min and then centrifuged at 3000 g for 15 min. Serum was collected and stored at -80℃ until further assay. The ELISA kit for corticosterone (QuicKey Pro Mouse CORT(Corticosterone) ELISA Kit, Elabscience) was used to assay the level of the hormones in serum^[71]^.

### Cortisol Measurement

Serum samples from MDD patients were collected as previous described and stored at -80℃ until further assay. The ELISA kit for cortisol (QuicKey Pro Human Cortisol ELISA Kit, Elabscience) was used to assay the cortisol levels in serum^[71]^.

### Surgical Procedure

Mice were anesthetized with sodium pentobarbital (40 mg/kg, intraperitoneally) and fixed in a stereotaxic head holder (Stoelting Co. , 51925). Eye ointment was applied to prevent eye dryness. The scalp was shaved, cleaned with distilled water, and sterilized with 75% ethanol, then a sagittal incision was made to expose the skull. The head was positioned to align the bregma and lambda landmarks by adjusting the head holder. A 0.5-mm hole was drilled on the top of the lateral ventricle (LV) using an electric drill (Hartmetall Instrumente, HM1005) until the surface of brain tissue was exposed, and remaining bone fragments were swept with fine forceps and cotton swabs^[72]^.

### Colchicine injections

A glass capillary (A-M Systems, 626000) was pulled into a micropipette with a neck length of 8–9LJmm and a tip diameter of 10 μm. Then a microsyringe (Hamilton, 701LJN) was attached to the micropipette and sealed with liquid paraffin. For colchicine injection, the microsyringe was loaded and fixed on the microinjection pump. Then 1μL of colchicine solution (6 µg/mL) was drawn into the microsyringe using the microinjection pump, and injected into the lateral ventricle (AP: ±1.0 mm, ML:

±0.5 mm, DV: -1.8 mm, based on the mouse brain atlas) at a rate of 50 nL/min. After the injection, the microsyringe was remained in place for 5-10 min before withdrawal to minimize backflow. Then the skull surface was cleaned, and the incision was stitched and steriled. Mice were sacrificed 48 h after injection, and brains were collected for CRH immunofluorescence staining^[73]^.

### Preparation of brain slices

Mice were anesthetized with 40 mg/kg 2% sodium pentobarbital by intraperitoneal injection and euthanized by cardiac perfusion. Blood was replaced with PBS, followed by perfusion with 4% paraformaldehyde for fixation. Then the whole brains were collected and fixed in 4% paraformaldehyde at 4°C for 48 hours, and dehydrated in 30% (w/v) sucrose PBS solution. The brains were embedded in OCT compound and quick-frozen at -20°C, and sliced into 50-μm-thin sections using a cryostat (Leica CM1950). The sections were stored in ethylene glycol antifreeze solution (PBS: ethylene glycol: glycerol = 50%: 30%: 20%(v/v)) at -20°C for following immunofluorescence staining.

### Immunofluorescence staining

The sections were washed 3 times with PBS, and were permeated (0.3% Triton X-100 (v/v) in PBS) for 30 min at 37°C. Then the sections were blocked in blocking solution (5% donkey serum (v/v), 0.3% Triton X-100 (v/v) in PBS) for 30 min at 37°C after washed with PBS. The primary antibodies were incubated for over 24h at 4°C; the sections were then washed 3 times with PBS and incubated with secondary antibodies at dilutions of 1:200 in PBS for 4 h at room temperature. After 3-time wash with PBS, all slices were mounted on slides and sealed with 80% glycerol for imaging. For sections stained with CRH neurons and adenosine A1 receptors, the sections were counterstained with DAPI in PBS for 10 min, then washed 3 times with PBS, and mounted on slides and sealed with 80% glycerol for imaging.

Primary antibodies included c-Fos (9F6) Rabbit mAb (Cell Signaling Technology, 2250S; 1:2000), Guinea Pig anti Corticotropin Releasing Factor (CRF) (human, mouse, rat) (BMA Biomedicals, T-5007; 1:800), Adenosine A1R Antibody - BSA Free (Novus Biologicals, NB300-549; 1:500) and Anti-PSD95 antibody [EPR23124-118] - Synaptic Marker (abcam, ab238135; 1:100). Secondary antibodies included Alexa Fluor® 488 AffiniPure® Donkey Anti-Rabbit IgG (H+L) (Jackson ImmunoResearch, 711-545-152), Alexa Fluor® 488 AffiniPure® Donkey Anti-Guinea Pig IgG (H+L) (Jackson ImmunoResearch, 706-545-148) and Alexa Fluor® 594 AffiniPure® Donkey Anti-Rabbit IgG (H+L) (Jackson ImmunoResearch, 711-585-152).

### Image acquisition

Three imaging strategies were implemented depending on imaging requirements. For imaging of c-Fos staining, the sections were visualized using TissueFaxs Plus (Tissue Genostics) imaging system. For imaging of CRH neurons and memory-related brain regions, the sections were visualized using Leica Stellaris 5 confocal microscope platform (Leica Microsystems). For imaging of colocalization of CRH neuron and adenosine A1 receptors, the sections were visualized using Zeiss LSM 980 confocal microscope (ZEISS Group).

## QUANTIFICATION AND STATISTICAL ANALYSIS

### Statistical analysis

The sample size chosen for our animal experiments in this study was estimated based on our prior experience of performing similar sets of experiments. All animal results were included, and no randomization method was applied. For all bar graphs, data were expressed as mean ± standard error of the mean (SEM). Statistical analysis was performed using unpaired Student’s t-tests for two groups and one-way analysis of variance (ANOVA) or two-way ANOVA (GraphPad Prism Software) for multiple groups, with all data points(https://www.graphpad.com/) showing a normal distribution. To compare two non-parametric datasets, a MannWhitney U-test was employed. p values <0.05 were considered significant. The sample sizes (biological replicates), specific statistical tests, and the main effects of the statistical details of experiments can be found in each figure legend.

## Data and materials availability

Data and materials are available from the corresponding author on reasonable request.

