## Supplementary Figures for "Gut Microbiota-derived Adenosine Determines the Efficacy of Electroconvulsive Therapy for Depression"

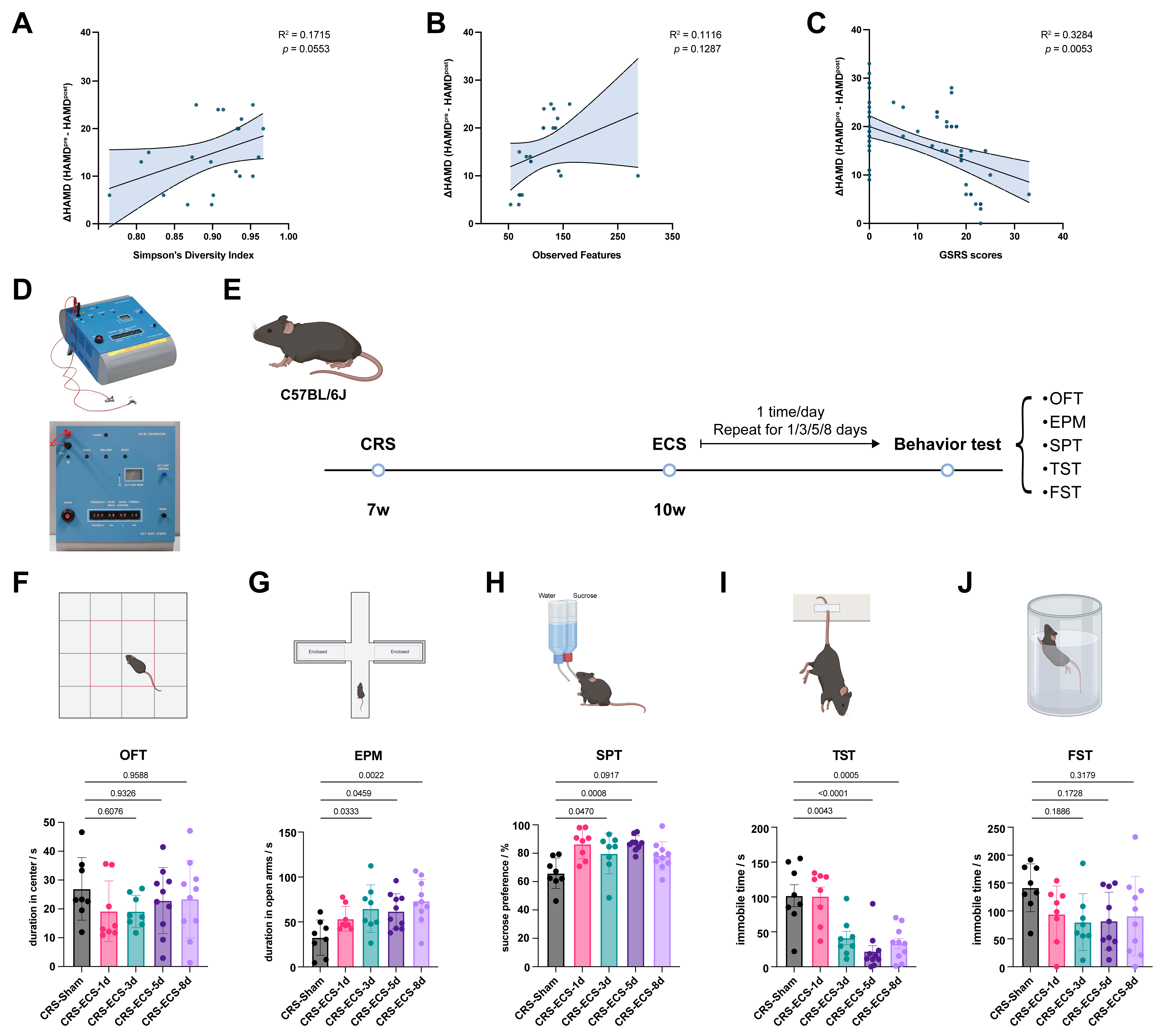

**Fig. S1 The improvement in antidepressant-like behavior peaked after a 5-day course of ECS.**

A-C. After receiving electroconvulsive therapy (ECT), the reduction in Hamilton Depression Rating Scale (HAMD) scores in patients with depression showed a tendency of positive correlation with the Simpson's diversity index (A) and observed features (B) in the alpha-diversity assessment and a significant negative correlation with GSRS scores (C).

A. ECS unit for mouse used in this study.
B. Timeline of different ECS treatment durations. A CRS model was established in mice aged 7 to 10 weeks. The mice were then treated by daily ECS for 1/3/5/8 days, with depression-related behavioral tests after the ECS treatment ended.
C. Duration into center area of mice in open-field test (OFT).
D. Duration into open arms of mice in elevated plus maze (EPM) test.
E. The preference for sucrose of mice in sucrose preference test (SPT).

I-J. The immobile time during the test of the mice in the tail suspension test (TST, I) and forced swimming test (FST, J).

Data are mean ± S.E.M., as determined by ordinary one-way ANOVA with Tukey correction, n = 8-10, the significance between groups is expressed numerically.

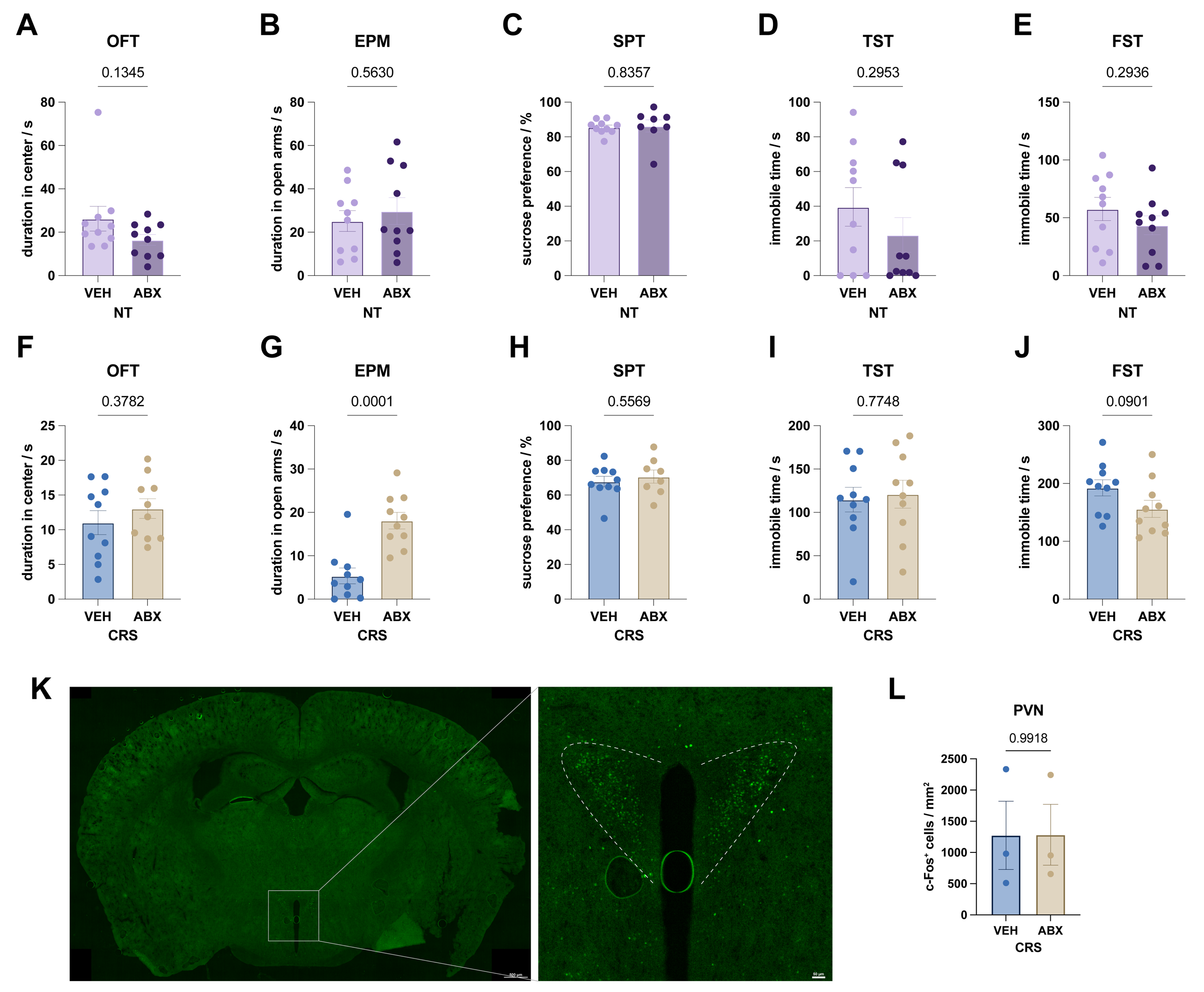

**Fig. S2 ABX treatment neither induced depressive-like behavior, nor altered the c-Fos activity of PVN.**

A. There was no significant difference in duration into center area of OFT between mice treated with vehicle and ABX.
B. There was no significant difference in duration into open arms of EPM between mice treated with vehicle and ABX.
C. There was no significant difference in preference for sucrose of SPT between mice treated with vehicle and ABX.

D-E. The immobile time during the test of the mice in the tail suspension test (TST, F) and forced swimming test (FST, G).

F. Duration into center area of mice in open-field test (OFT).
G. Duration into open arms of mice in elevated plus maze (EPM) test.
H. The preference for sucrose of mice in sucrose preference test (SPT).

I-J. The immobile time during the test of the mice in the tail suspension test (TST, F) and forced swimming test (FST, G).

K. Representative c-Fos imaging of PVN of CRS-ABX-Sham mice.
L. The numbers of c-Fos-expressing neurons per mm^2^ in PVN of CRS-ABX-Sham mice.

Data are mean ± SEM, as determined by ordinary one-way ANOVA with Tukey correction, the significance between groups is expressed numerically, ns = no significance.

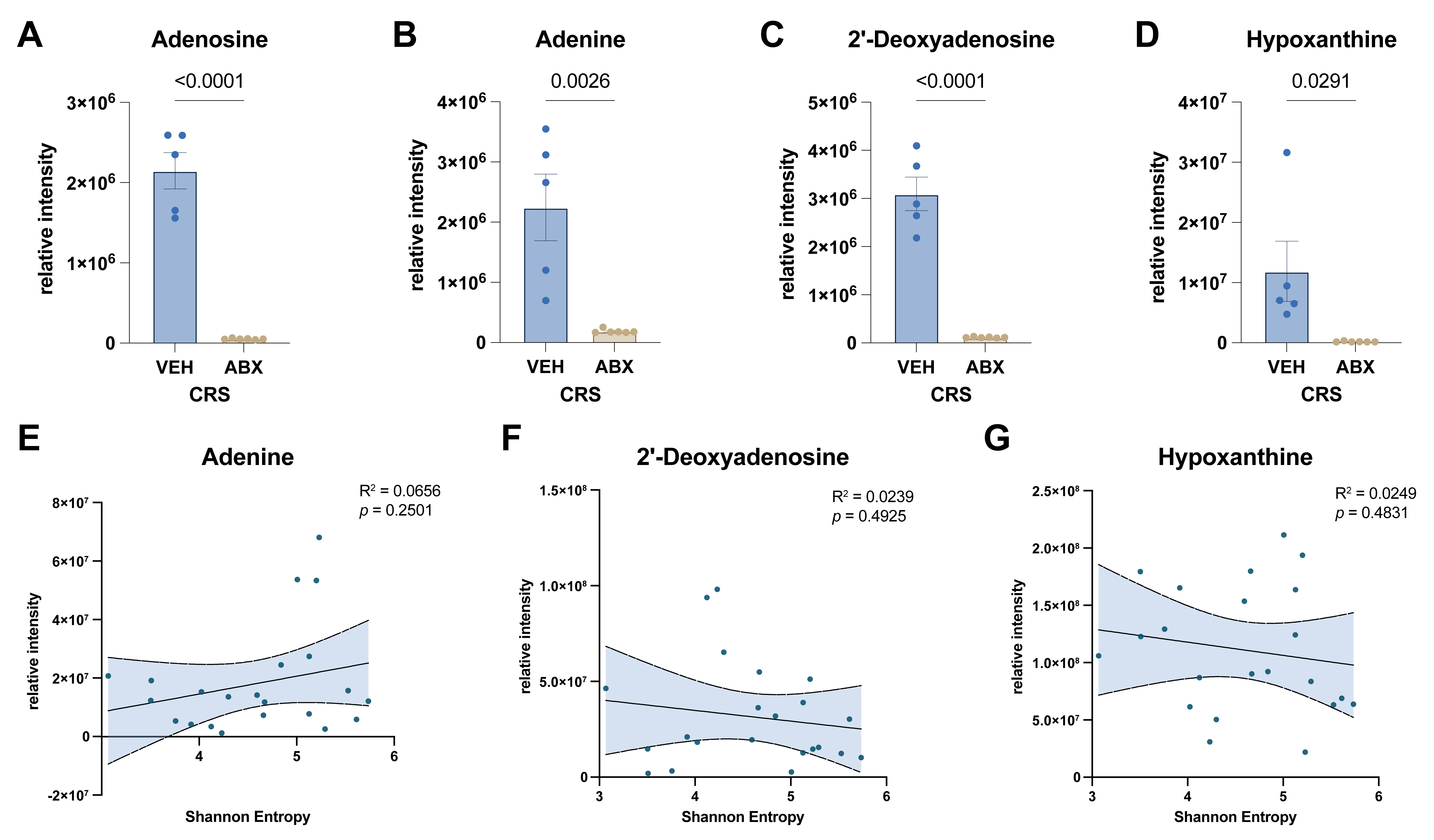

**Fig. S3 ABX treatment reduced the amount of metabolites in the purine metabolism pathway.**

A-D. The relative intensity of adenosine (A), adenine (B), 2’-deoxyadenosine (C) and hypoxanthine (D) in feces of CRS-ABX-ECS- and CRS-VEH-ECS mice.

E-G. The correlation analysis between the relative intensity of adenine (E), 2’-deoxyadenosine (F) and hypoxanthine (G) in feces of MDD patients and their Shannon entropy.

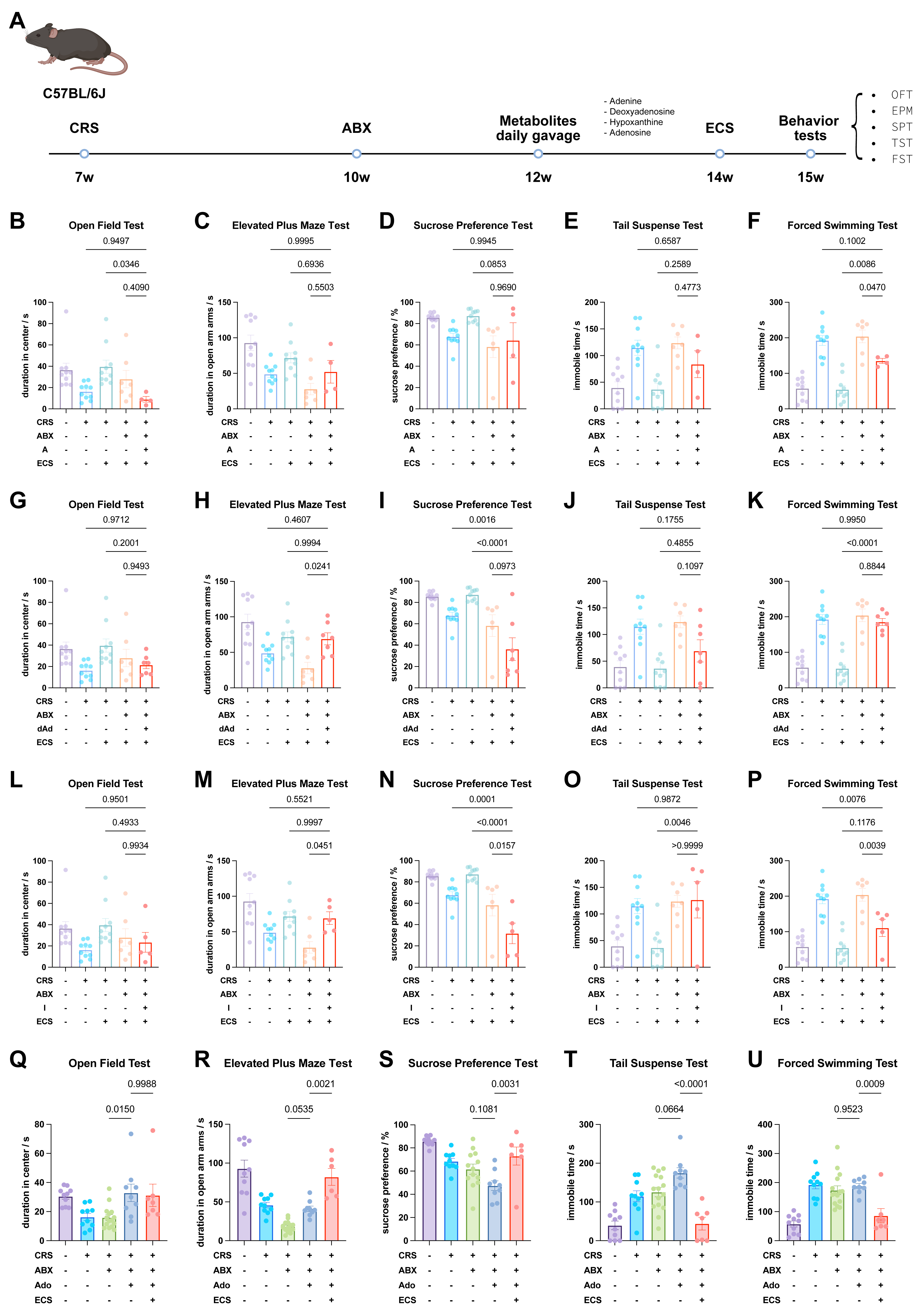

**Fig. S4 Supplement of adenine, 2’-deoxyadenosine and hypoxanthine to ABX-treated depressant mice showed no significant recoveries in antidepressant effects of ECS**

A. Timeline of the behavioral tests on metabolite replenishment. A CRS model was established in mice aged 7 to 10 weeks. The mice were then treated with ABX or vehicle for 2 weeks, followed by daily gavage of metabolites supplements for 2 weeks, and finally underwent ECS or sham surgery for 5 days. Behavioral tests were performed on the mice at 15 weeks of age, and tissue samples were collected after all behavioral tests were completed.

B-F. Duration in center of the OFT (B), duration in open arm of the EPM (C), sucrose preference of the SPT (D), immobile time in TST (E) and (FST) of mice gavaged by adenine.

G-K. Duration in center of the OFT (B), duration in open arm of the EPM (C), sucrose preference of the SPT (D), immobile time in TST (E) and (FST) of mice gavaged by 2’-deoxyadenosine.

L-P. Duration in center of the OFT (B), duration in open arm of the EPM (C), sucrose preference of the SPT (D), immobile time in TST (E) and (FST) of mice gavaged by hypoxanthine.

Q-U. Duration in center of the OFT (B), duration in open arm of the EPM (C), sucrose preference of the SPT (D), immobile time in TST (E) and (FST) of mice gavaged by adenosine.

Data are mean ± SEM, as determined by ordinary one-way ANOVA with Tukey correction, the significance between groups is expressed numerically.

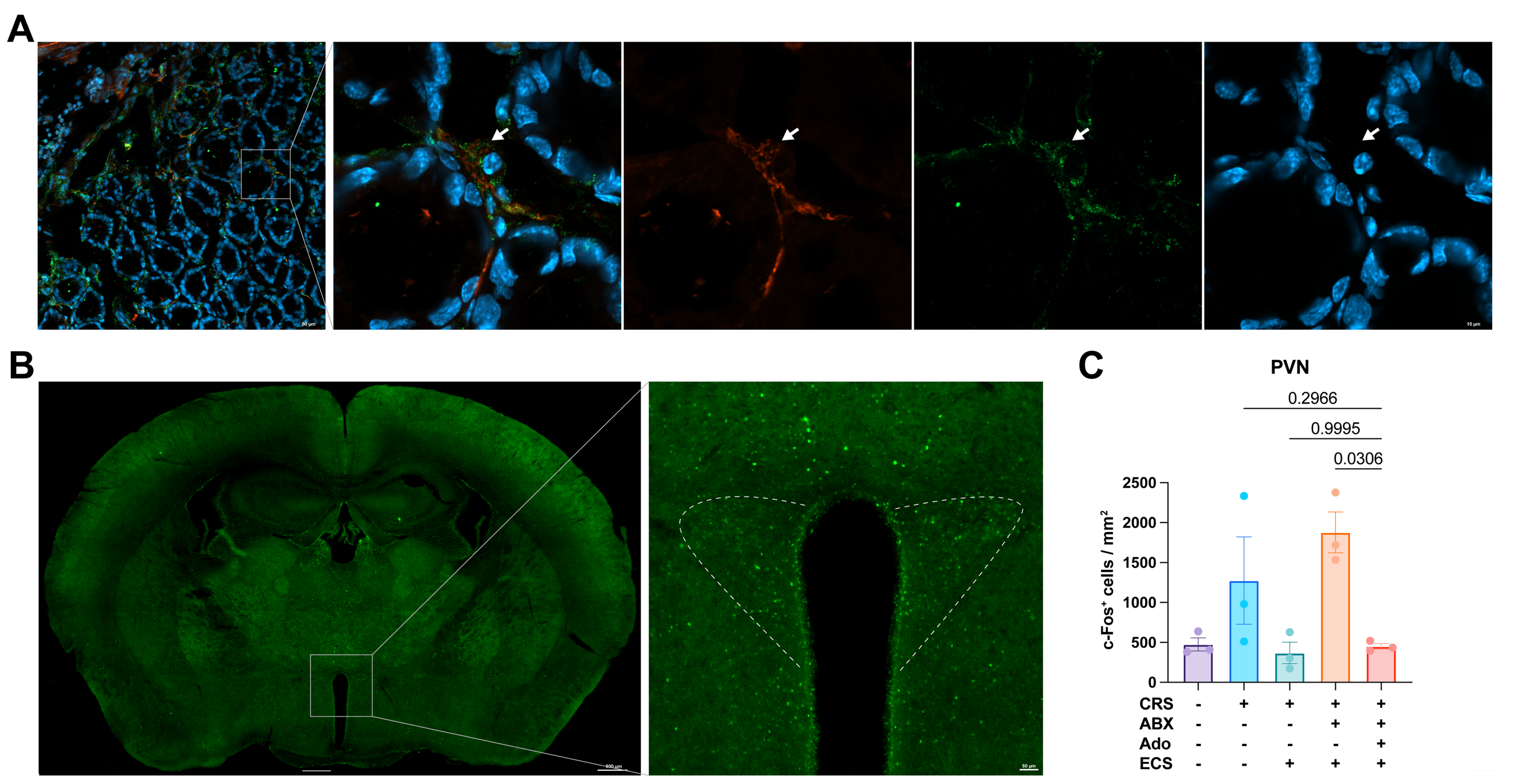

**Fig. S5 Adenosine supplement to ABX-treated depressant mice recovered down-regulation of PVN c-Fos activity after ECS.**

A. Representative c-Fos imaging of PVN of CRS-ABX-Adenosine-ECS mice.
B. The numbers of c-Fos-expressing neurons per mm2 in PVN of CRS-ABX-Adenosine-ECS mice. Data are mean ± SEM, as determined by ordinary one-way ANOVA with Tukey correction, the significance between groups is expressed numerically.
C. Representative imaging showing the connection between adenosine receptor A1(Adora1) expressing-intestinal epithelium cell and UCHL1-labeled enteric neuron.

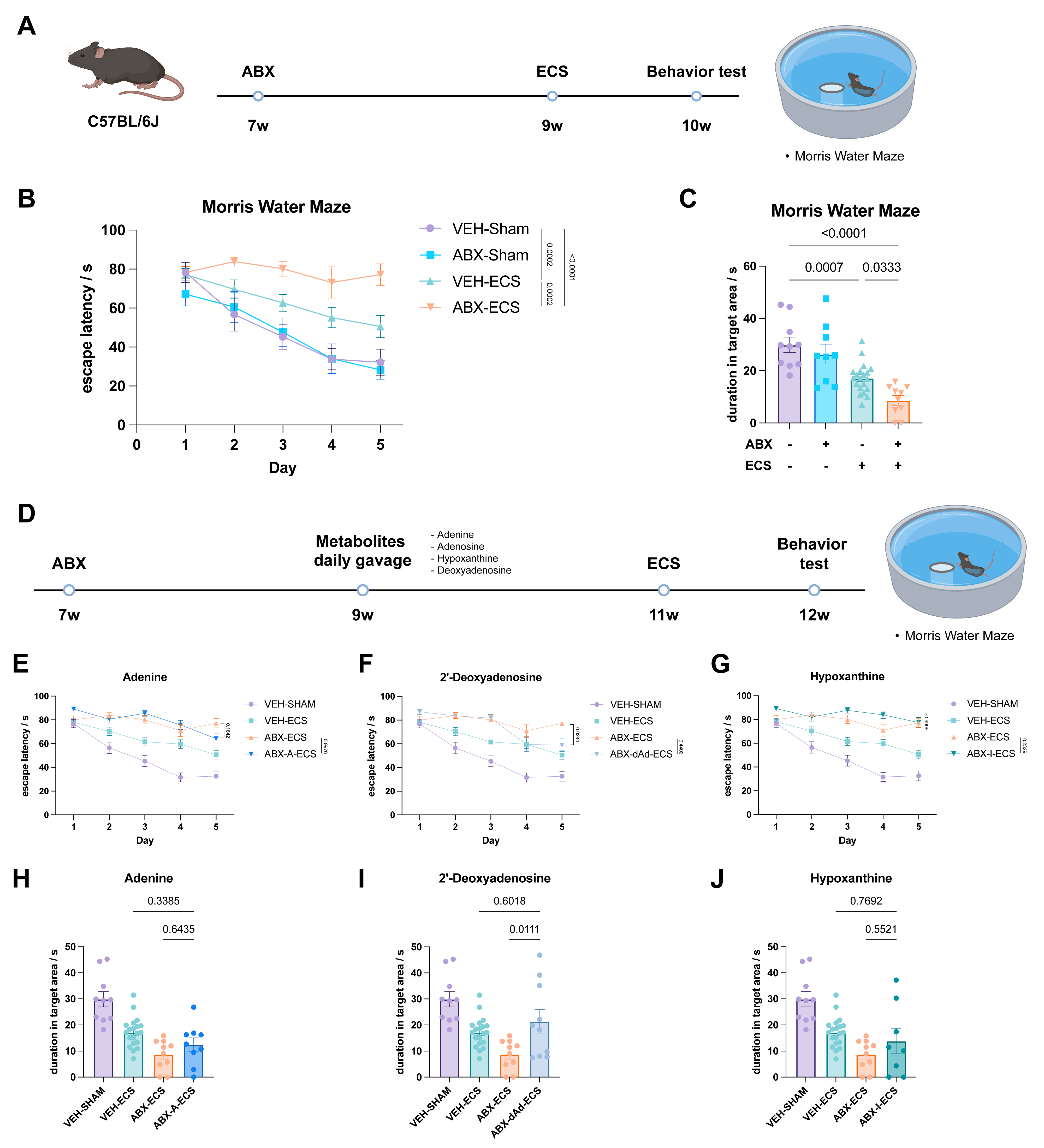

**Fig. S6 Adenine or hypoxanthine supplement to ABX-treated mice did not ameliorate the impairment of spatial memory after ECS.**

A. Timeline of experiments on ABX-treatment impairment of spatial memory after ECS. The mice were treated with ABX or vehicle for 2 weeks, followed by ECS or sham surgery for 5 days. Morris water maze (MWM) test was performed on the mice at 10 weeks of age.
B. Escape latency in acquisition period of all groups of mice by day.
C. Time lingering in target area of all groups of mice at the last day of testing.
D. Timeline of experiments on metabolite replenishment to alleviate memory loss. The mice were treated with ABX or vehicle for 2 weeks, followed by daily gavage of metabolites supplements for 2 weeks, and finally underwent ECS or sham surgery for 5 days. MWM tests were performed on the mice at 12 weeks of age.

E-G. Escape latency in acquisition period of ABX-treated mice supplemented with 2’-deoxyadenosine (E), adenine (F) and hypoxanthine (G) by day.

H-J. Time lingering in target area of ABX-treated mice supplemented with 2’-deoxyadenosine (H), adenine (I) and hypoxanthine (J) at the last day of testing.

Data are mean ± SEM, as determined by ordinary two-way ANOVA with Tukey correction, the significance between groups is expressed numerically.
